# Chemically diverse antagonists inhibit human TRPV1 through ligand-specific interaction networks

**DOI:** 10.64898/2026.04.27.721197

**Authors:** Kyle E. Lopez, Audrey S. Paduda, Matthew J. Derrick, Wade D. Van Horn

## Abstract

Transient receptor potential vanilloid 1 (TRPV1) mediates thermosensation and pain and is a target for non-addictive analgesics; however, clinical candidates have failed due to thermoregulatory and thermosensory side effects. Limited structural data for human TRPV1 (hTRPV1) bound to clinically relevant antagonists has constrained mechanistic insight. Using chemoinformatics-informed cryo-EM and BRET assays, we identify variable antagonist target engagement across diverse chemotypes, including failed clinical compounds. A structure of hTRPV1 bound to 6-iodo-dihydrocapsaicin shows how a single atom substitution converts an agonist into an antagonist. Additional structures with Asivatrep, Mavatrep, and JNJ-17203212 reveal vanilloid pocket plasticity and divergent interaction networks, including lipid co-binding. Mutagenesis of ligand-interacting residues further demonstrates distinct contributions to antagonist engagement. Despite this diversity, antagonists converge on a conserved inhibited state, showing high potency is maintained across flexible binding modes. These findings refine our understanding of hTRPV1 antagonism and inform strategies for next-generation analgesics with improved safety.

## Introduction

Chronic pain is a pervasive global health problem^1^. In the United States, chronic pain is estimated to affect 20.9% of adults (∼51.6 million people), and opioids remain among the most effective analgesics available^2^. Unfortunately, opioids are associated with unwanted risks such as addiction and overdose, which has prompted deep interest in alternative nociceptive targets without the liabilities of opioid treatment^3, 4, 5^. Recent clinical success with ion channel-targeted analgesics, such as the Na_V_1.8 antagonist Journavx, underscores the therapeutic potential of non-addictive ion channel inhibitors^6, 7, 8^. One such target is the transient receptor potential vanilloid type 1 (TRPV1), a polymodally activated and allosterically regulated, non-selective cation channel that senses noxious heat, chemical stimuli, and inflammatory signals and plays a central role in thermoregulation, thermosensing, and pain sensation^9, 10, 11, 12, 13, 14, 15, 16^.

Early pharmacological efforts to target TRPV1 for pain relief focused on agonists such as capsaicin and resiniferatoxin (RTX), but the intense burning sensation associated with channel activation has generally limited agonist use to topical applications^17^. The prototypical TRPV1 agonist capsaicin, the pungent component from chili peppers, remains the only FDA-approved TRPV1 agonist and is used in a topical cream to treat arthritic pain and an 8% capsaicin patch (Qutenza) to treat shingles pain and diabetic peripheral neuropathy^18, 19^. These challenges shifted the therapeutic strategy, and most subsequent drug development efforts have focused on TRPV1 antagonism^10, 20, 21, 22^.

While TRPV1 antagonism is generally considered the more efficient route to desired clinical outcomes, TRPV1-specific antagonists have failed clinical trials due to on-target adverse effects, most notably hyperthermia and thermohypoesthesia (impaired heat sensation), though the extent of these effects is antagonist dependent^17, 22, 23, 24^. These outcomes highlight a critical gap in understanding how different antagonists engage TRPV1 and modulate distinct activation modes^14^. Clinical and preclinical data suggest that mode-selective TRPV1 antagonists are key to avoiding unwanted thermoregulatory and thermosensory side effects. Complicating TRPV1 drug discovery is that rodent and human TRPV1 differ in pH sensitivity and antagonist response, and promising compounds in preclinical studies often fail to translate to the clinic^24, 25, 26^. These species-specific differences limit the predictive power of preclinical studies and highlight the need to study the human TRPV1 ortholog specifically to define the molecular mechanisms for antagonism. Notably, human genetic evidence demonstrates that TRPV1 directly regulates pain sensitivity, as complete loss-of-function mutations alter nociception and abolish capsaicin-evoked pain without eliminating other pain modalities^12^. Similarly, naturally occurring human TRPV1 variants can selectively reduce nociceptive signaling and injury-induced hypersensitivity when transplanted to rodents while preserving normal heat sensation and thermoregulation^13^. Together, these data validate TRPV1 as a genetically supported pain target and demonstrate that modality-selective modulation can potentially provide efficacious analgesia without thermoregulatory or thermosensory liabilities. At the molecular level, ligand binding within the orthosteric vanilloid pocket is canonically anchored by interaction with Y511, while functional outcomes are coupled to gating through allosteric communication between the voltage-sensing like domain (VSLD) and pore via the S4-S5 linker, generally involving the R557-E570 interaction^27, 28, 29, 30, 31, 32, 33, 34^. Despite conservation of these residues and channel architecture, vanilloid sensitivity has diverged across species, most notably birds, whose TRPV1 channels exhibit markedly reduced sensitivity to capsaicin, consistent with the absence of capsaicin pungency in birds^14, 15, 35, 36^. Y511 and the neighboring S512 are conserved across vertebrate TRPV1 orthologs and contribute directly to vanilloid recognition and sensitivity^35, 37^. Thus, conservation of these key ligand-interacting residues does not necessarily correspond to conserved vanilloid sensitivity and highlights how differences in interaction networks and allosteric pathways can produce substantial changes in pharmacological response. Agonists and antagonists differentially modulate these interactions, altering the S4-S5 linker position and regulating channel gating (Fig. 1a)^27, 28, 29, 30, 31, 32, 33, 34^. Consistent with this, a potentially informative molecule to understand TRPV1 antagonism is 6-iodo-dihydrocapsaicin (6-Iodo-CAP, CAY10448), in which an iodine moiety is added at the 6ʹ position in dihydrocapsaicin. This single-atom change that replaces a hydrogen with an iodine, converts the canonical agonist into a potent antagonist^38^. Because 6-Iodo-CAP preserves the vanilloid scaffold, it is a useful tool to probe the minimum requirements for competitive channel inhibition. This minimal conversion to antagonist establishes a reference point for comparison with more diverse TRPV1 antagonist chemotypes that achieve inhibition through distinct interactions.

**Fig. 1:**
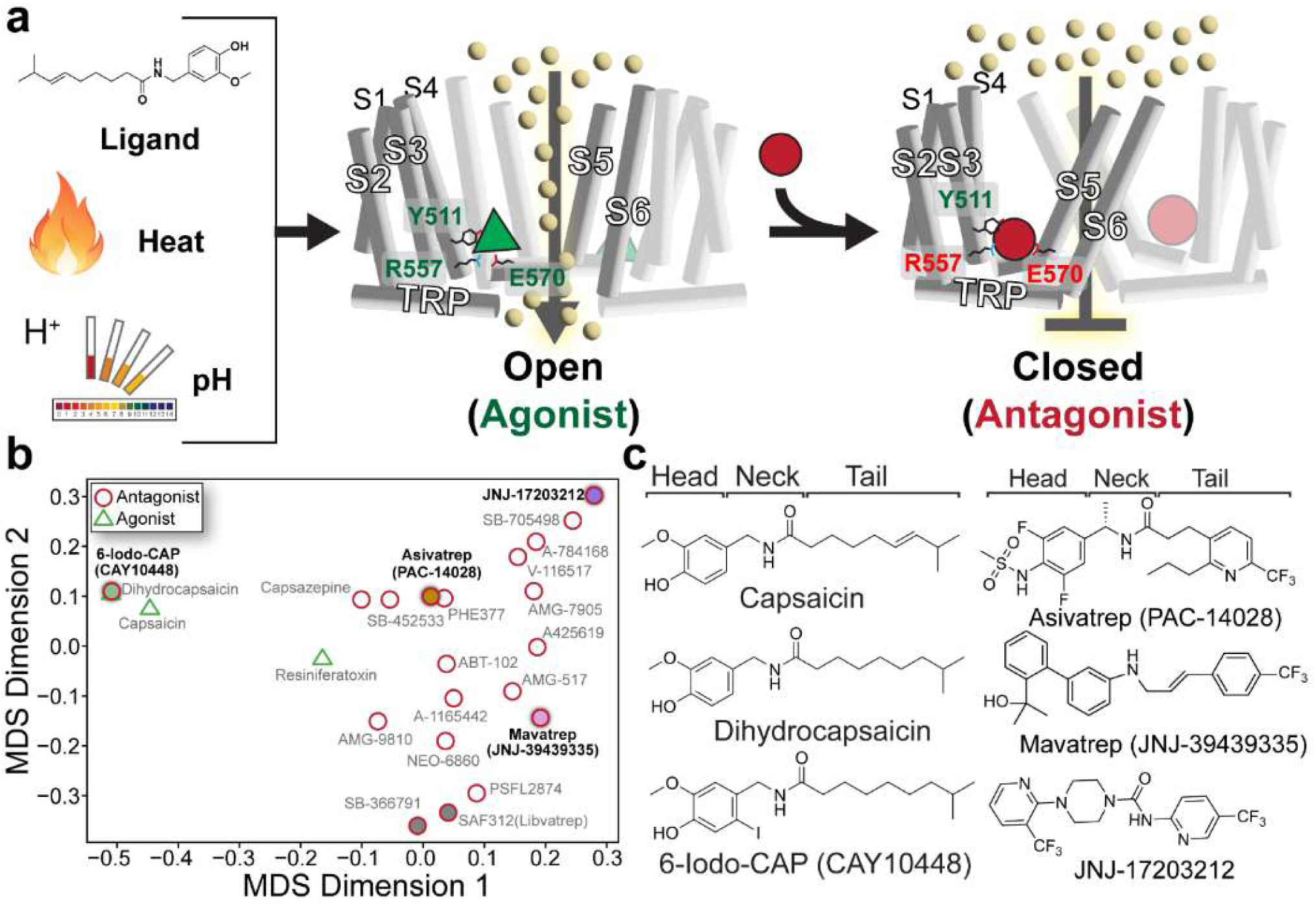
TRPV1 agonist-antagonist gating model and scaffold diversity across ligand chemotypes. **a** Schematic of TRPV1 gating that illustrates the transition from an open, agonist-bound state (middle, green) to a closed, antagonist-bound state (right, red). Agonist stimuli (ligand, heat, and protons) are shown to the left. Key residue interactions are labeled in green and interaction disruptions in red. **b** Chemoinformatic latent space representation of selected ligands that maps and quantifies TRPV1 chemotype diversity. Agonists are labeled with green triangles and antagonists with red circles. Circles filled grey (bottom right) represent published hTRPV1 structures bound to specified antagonists; other colored circles (green, orange, pink, purple) represent the antagonist structures that are investigated in this study. **c** Chemical structures for Capsaicin, Dihydrocapsaicin, and 6-Iodo-CAP (left) and Asivatrep, Mavatrep, and JNJ-17203212 (right) with head, neck, and tail regions labeled above.

To further understand hTRPV1 antagonism, we examine a set of chemically diverse, clinically relevant TRPV1 antagonists. The antagonist JNJ-39439335 (Mavatrep) was involved in 5 clinical studies to treat osteoarthritis and demonstrated analgesic efficacy, but was terminated after a phase I study due to hyperthermia and thermohypoesthesia^39, 40, 41, 42, 43, 44^. Recently, MAC Clinical Research announced plans to reinitiate phase 2 clinical trials for Mavatrep targeting chronic osteoarthritis pain. PAC-14028 (Asivatrep) progressed through phase 3 clinical trials in Korea to treat atopic dermatitis and is currently used in a cosmetic cream (AESTURA) that treats dry, itchy skin^45, 46, 47, 48, 49, 50^. Lastly, the preclinical antagonist JNJ-17203212 showed efficacy across multiple rodent pain models that include migraine, osteoarthritis, and colonic hypersensitivity, and is among the most chemically distinct TRPV1 antagonist chemotypes^51, 52, 53, 54, 55^.

Despite the known differences between human and other TRPV1 orthologs (vide supra), the approaching 100 TRPV1 clinical trials listed in clinicaltrials.gov, and a large number (>80) of TRPV1 structures in the RCSB Protein Structure Database; there is a paucity of human TRPV1 structural information ^27, 28, 29, 30, 31, 32, 33, 34, 56, 57, 58, 59, 60^. To date, there are 9 total published hTRPV1 structures and only two human TRPV1 (hTRPV1) structures with an antagonist bound to the orthosteric vanilloid binding site, where clinically relevant antagonist drug discovery has focused, have been published. One additional hTRPV1 structural study identifies allosteric antagonism from a non-selective natural product^57^. To address this gap, we performed chemoinformatic analysis to map the TRPV1 chemotype space for antagonists that bind the orthosteric vanilloid binding site using a multidimensional scaling (MDS) approach (Fig. 1b, Supplementary Table 1). From this quantitative analysis, we selected a set of chemically diverse antagonists that include 6-Iodo-CAP, a minimal agonist-to-antagonist transition, alongside the clinically relevant compounds Mavatrep, Asivatrep, and JNJ-17203212 (Fig. 1c). We combine high-resolution cryo-electron microscopy (cryo-EM) structures and functional bioluminescence resonance energy transfer (BRET) assays coupled with targeted mutagenesis to investigate how these distinct chemotypes engage hTRPV1. Our results reveal how chemically diverse ligands exploit conformational plasticity within a shared vanilloid binding pocket to stabilize a common inhibited state. These findings redefine principles for TRPV1 ligand recognition and orthosteric modulation in TRP channels and provide a framework for future structure-guided design of TRPV1 ion channel therapeutics.

## Results and Discussion

### Cryo-EM structure of hTRPV1 solubilized in detergent retains endogenous lipids

Full-length human TRPV1 was expressed in HEK293 GnTI^-^ cells, purified in glycol-diosgenin (GDN) detergent, and used for cryo-EM studies. To assess whether detergent solubilization preserves a physiologically relevant conformation and lipid environment, we determined an apo hTRPV1 (hTRPV1_Apo_) structure in GDN (Fig. 2a, b, Supplementary Fig. 1, Supplementary Table 2). The structure was resolved to 2.5 Å resolution, comparable to or higher than previously reported nanodisc-reconstituted apo hTRPV1 structures (2.9 Å, PDB 8GF8; 2.6 Å, PDB 8GF9)^29^.

**Fig. 2:**
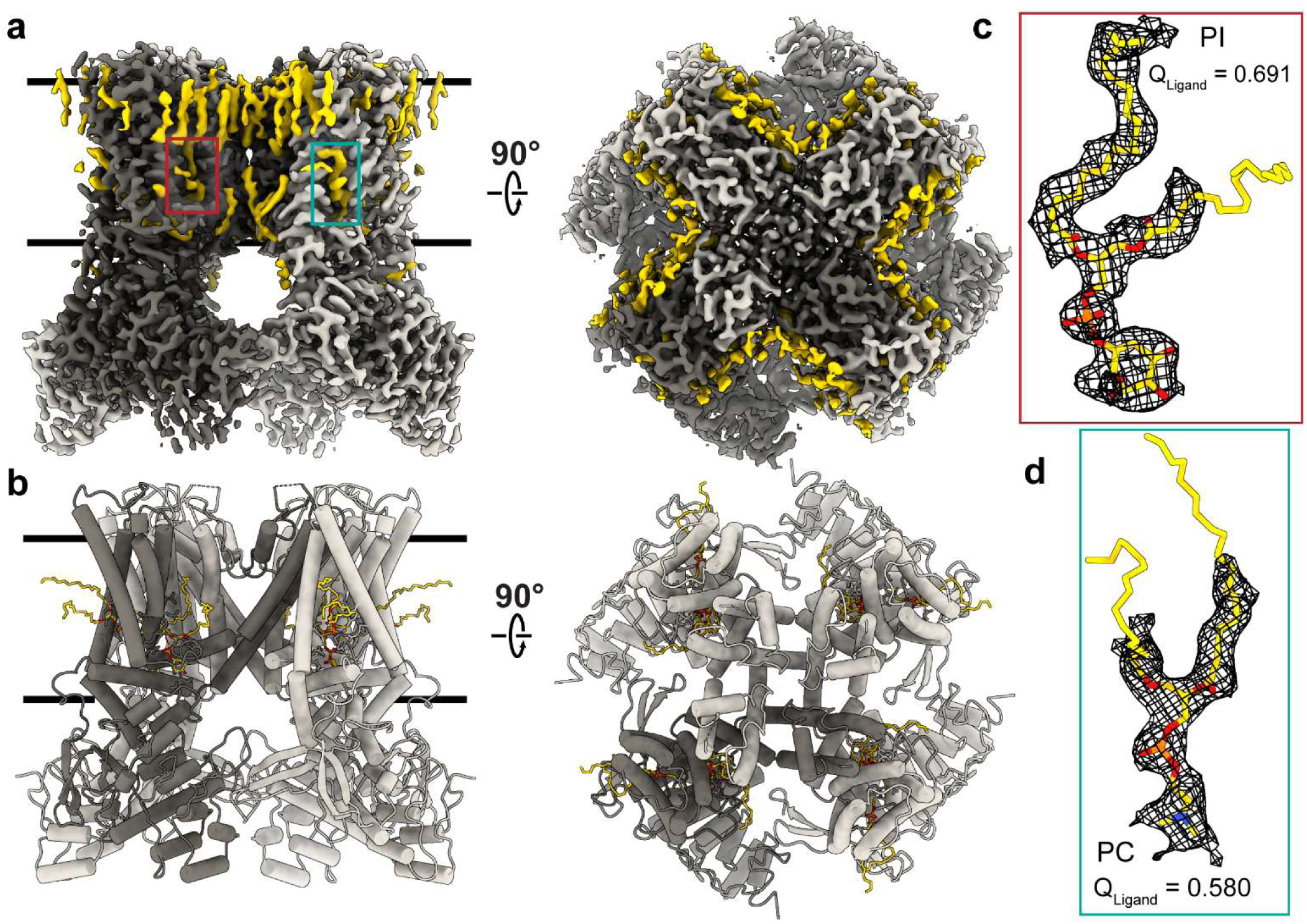
Apo human TRPV1 structure in GDN retains bound lipid densities. **a** Cryo-EM density for hTRPV1_Apo_ in GDN, colored grey with a monomer colored a dark grey and lipid densities in yellow. **b** hTRPV1_Apo_ atomic model shown with the same color scheme as **a**. Cryo-EM density for phosphatidylinositol (PI) (**c**) and phosphatidylcholine (PC) (**d**) shown as black mesh with lipids shown in yellow. Corresponding ligand Q-scores (Q_Ligand_) are indicated.

Consistent with previous nanodisc structures, Cryo-EM density from our GDN-solubilized hTRPV1 reveals the presence of many annular lipids that have been copurified with the channel (Fig. 2a-d).

Specifically, a phosphatidylinositol lipid (PI) within the vanilloid pocket and a phosphatidylcholine lipid (PC) at an adjacent site are identified (Fig. 2a-d). The expected PI and PC lipid densities are well resolved (Q_Ligand_ = 0.691 for PI; Q_Ligand_ = 0.580 for PC), which supports confident ligand model placement (Fig. 2c, d). Furthermore, the annular lipid densities in hTRPV1_Apo_ are inconsistent with GDN detergent, as they are too narrow to be the GDN steroidal hydrophobic tail. The apo structure adopts a closed conformation and closely matches nanodisc-reconstituted hTRPV1, with a similar domain organization in the transmembrane domain and intracellular ankyrin repeats. Structural alignment yields Cα root mean squared deviation (RMSD) of 1.03 Å relative to PDB 8GF8 and 1.02 Å relative to PDB 8GF9, which corresponds to a normalized RMSD_100_ of 0.33 and 0.31, respectively. Together, these results indicate that hTRPV1 purified in GDN retains key structural and environmental features, such as lipid interactions, observed in nanodisc systems, supporting the use of detergent-solubilized hTRPV1 for comparative structural analysis of antagonist-bound states.

### How a single atom substitution converts the common agonist dihydrocapsaicin into a TRPV1 antagonist

Capsaicin and the common hydrogenated tail variant dihydrocapsaicin (Fig. 1b) which comprises ∼30-50% of total capsaicinoids found in common chili peppers, are prototypical TRPV1 agonists that bind the orthosteric vanilloid binding pocket^61^. To establish a structural reference for agonist recognition by the human channel, we determined a cryo-EM structure of hTRPV1 bound to capsaicin (hTRPV1_CAP_) at 3.06 Å resolution (Fig. 3a, Supplementary Fig. 2). Previous capsaicin-bound TRPV1 structures were determined at lower resolutions, and none used the human ortholog^30, 33, 58^. At 3.06 Å resolution, capsaicin can be unambiguously modeled into the ligand density (Q_Ligand_ = 0.792), allowing the interactions within the vanilloid binding pocket to be clearly resolved (Fig. 3b,c). Similar to the only other hTRPV1 structure bound to an agonist (MSP20) and other agonist-bound rodent structures, hTRPV1_CAP_ is in a non-conducting conformation (Fig. 3a, Supplementary Fig. 3)^62^. This is likely due to the rapid desensitization of TRPV1 followed by agonist binding that causes the channel to close. When hTRPV1_CAP_ is overlayed with rat TRPV1_Open_ (PDB 5IRX) by the transmembrane domain (residues 416 to 712), the RMSD is 1.19 Å with a majority of the differences occurring at the S6 and pore helices (Supplementary Fig. 3a, b). When two rat TRPV1 open structures are compared (5IRX and 7RQW) there are little to no differences in the S4–S5 linker and S6 regions with a 0.80 Å RMSD (Supplementary Fig. 3f).

**Fig. 3:**
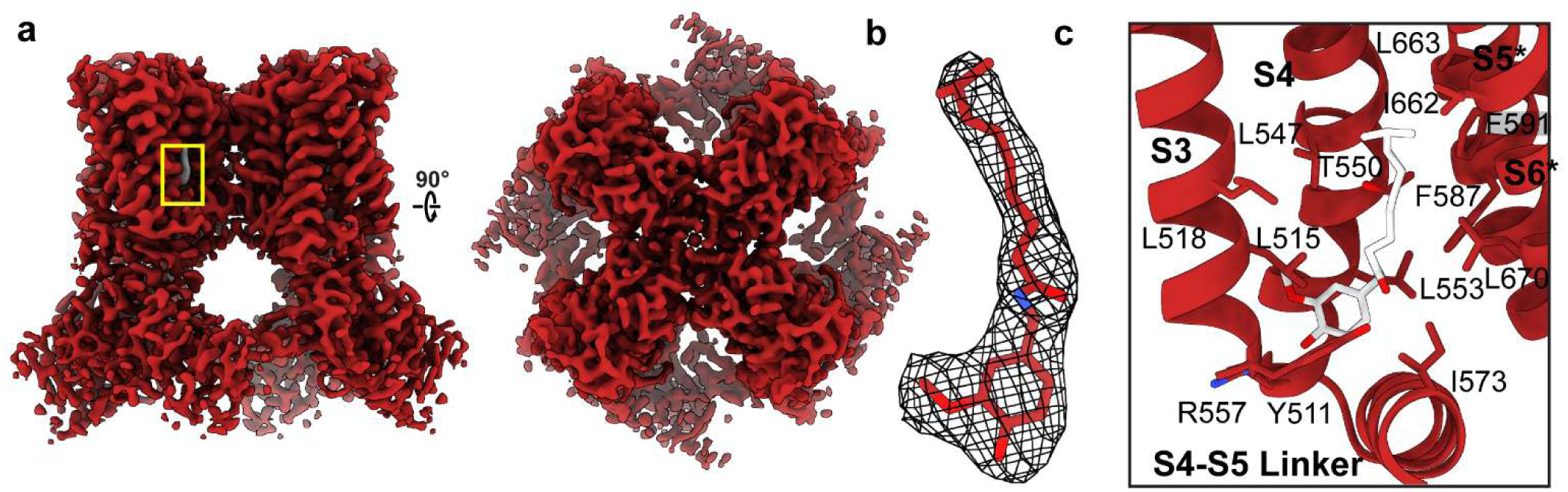
hTRPV1 bound to canonical agonist capsaicin is in a closed state. **a)** Cryo-EM density of hTRPV1_CAP_ side view (left) and top view (right) with hTRPV1 colored in red and capsaicin colored grey and vanilloid binding pocket outlined with a yellow box. **b)** Capsaicin modeled into cryo-EM density (black mesh; Q_Ligand_ = 0.792). **c)** Vanilloid binding pocket with capsaicin (white) and interacting residues represented as sticks.

Capsaicin and dihydrocapsaicin are both potent agonists, despite the difference in tail hydrogenation, and a Ca^2+^-sensing BRET assay-obtained *EC*_50_ values for capsaicin (p*EC*_50_ = 8.34 ± 0.06; *EC*_50_ *≈* 5 nM) and dihydrocapsaicin (p*EC*_50_ = 9.0 ± 0.1; *EC*_50_ *≈* 1.0 nM) suggest that double bond deletion plays a role in dihydrocapsaicinimpacts potency (Fig. 4a, b). The nearly five-fold enhancement in apparent potency observed for dihydrocapsaicin compared to capsaicin suggests insight into the structural plasticity of the orthosteric binding pocket. The sole chemical difference between these ligands is the *trans*-double bond in capsaicin’s aliphatic tail, a feature that limits the ligand conformational ensemble. If the orthosteric vanilloid binding site functioned as a static, rigid cleft contoured specifically to capsaicin, it would be expected to exhibit equal or increased affinity. Instead, the saturated, flexible tail of dihydrocapsaicin results in a significant apparent energetic increase in affinity of -0.9 ± 0.2 kcal/mol as calculated from a modified Cheng-Prusoff framework (see Methods)^63,64^. This thermodynamic preference for a conformationally unconstrained ligand suggests that the binding pocket is inherently dynamic. The site appears capable of local structural rearrangements that allow it to pack more efficiently around the flexible aliphatic chain, thereby optimizing hydrophobic contacts to stabilize the active state.

**Fig. 4:**
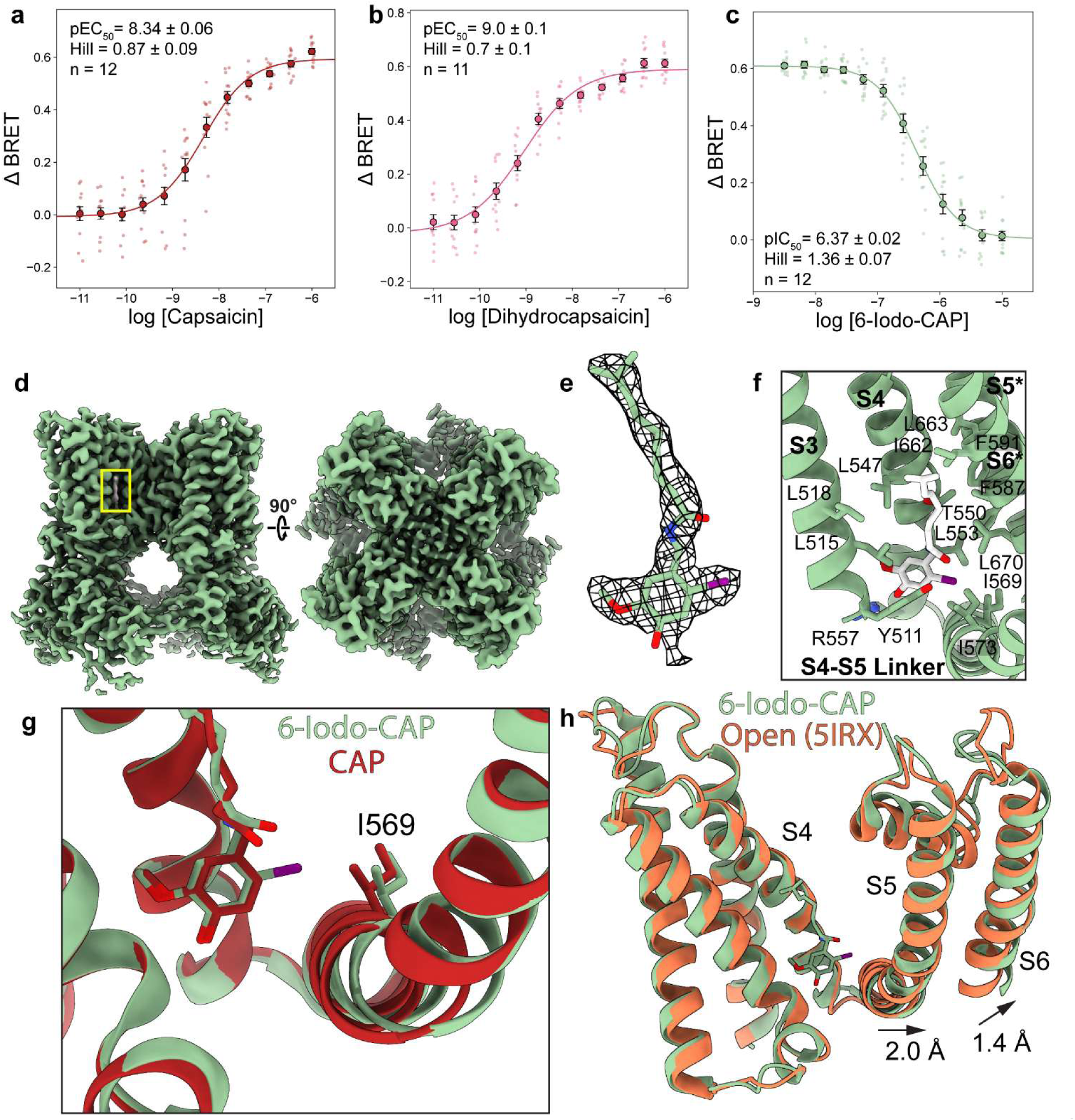
6-Iodo-CAP binding displaces the S4-S5 linker to stabilize the inhibited hTRPV1 state. BRET Ca^2+^ influx dose-response curves for capsaicin (**a**), dihydrocapsaicin (**b**), and 6-iodo-CAP in the presence of 250 nM capsaicin (**c**). **d** Cryo-EM density for hTRPV1_6-Iodo-CAP_ (green) side (left) and top (right) views; 6-Iodo-CAP density colored grey and location indicated with yellow box. **e** 6-Iodo-CAP cryo-EM density and stick representation 6-Iodo-CAP modeled into density (Q_Ligand_ = 0.784). **f** Vanilloid binding pocket with 6-Iodo-CAP with interacting residues shown as sticks and labeled. **g** Overlay between hTRPV1_CAP_ (red) and hTRPV1_6-Iodo-CAP_ (green) and I569 represented as sticks. Structures are aligned by the backbone residues for the transmembrane domain (416-712). **h** Overlay of hTRPV1_6-Iodo-CAP_ (green) and rTRPV1_Open_ (coral, PDB 5IRX) and arrows to indicate hTRPV1_6-Iodo-CAP_ helix displacement in hTRPV1_6-Iodo-CAP_ relative to rTRPV1_Open_. Structures are overlayed same as (**g**). For the BRET data in panels **a**-**c**, the jittered data represent distinct replicates with the dose average values shown as a colored circle; the error bars are SEM. The p*IC*_50_ and n_H_ error values are derived from fitting errors.

Previous studies show that the addition of a halogen at the 5ʹ or 6ʹ position in the vanilloid head group converts TRPV1 vanilloid agonists into antagonists (Fig. 1b)^38, 65^. Halogenation at the 6ʹ position results in a more potent antagonist relative to the 5ʹ position for dihydrocapsaicin analogs, but the opposite is true for halogenated RTX analogs^38, 65^. Furthermore, larger halogens result in more potent antagonists, with an iodine moiety resulting in the most potent antagoinism^38^. Our BRET assay confirms that 6-Iodo-CAP inhibits CAP stimulated Ca^2+^ flux and competes for the vanilloid pocket with capsaicin with a p*IC*_50_ of 6.37±0.02 (∼430 nM, Fig. 4c). The transformation of dihydrocapsaicin from a potent agonist into the antagonist 6-iodo-dihydrocapsaicin highlights the local spatial sensitivity of the vanilloid binding site during TRPV1 allosteric activation mechanism. When corrected for the high-agonist CAP competitive assay concentration (250 nM), 6-iodo-dihydrocapsaicin binds with single-digit nanomolar affinity for the resting state (*Ki* = 7.66 nM). However, this iodo halogenation decreases the apparent affinity with energetic penalty (ΔΔG) of +1.21 ± 0.16 kcal/mol relative to apparent agonist affinity. This thermodynamic shift, coupled with the complete loss of efficacy, indicates that the vanillyl 6-position occupies a critical mechanistic hinge within the orthosteric pocket. In a conformationally dynamic binding site, the resting-state pocket can readily accommodate the steric bulk and polarizability of the iodine atom. Yet, halogenation likely creates a severe steric clash inconsistent with activation.

Consequently, the iodine moiety presumably acts as a molecular wedge that permits high-affinity binding while physically obstructing the dynamic allosteric conformational propagation required to stabilize the open state.

To further investigate 6-Iodo-CAP antagonism, we determined the first hTRPV1 cryo-EM structure bound to 6-Iodo-CAP (hTRPV1_6-Iodo-CAP_) at 2.90 Å resolution (Fig. 4d, Supplementary Fig. 4, Supplementary Table 2). The local resolution in the vanilloid binding pocket is even higher (∼2.4 Å) and the corresponding 6-Iodo-CAP density allows the compound to be modeled unambiguously into the density (Ligand Q-score (Q_Ligand_) = 0.784, Fig. 4e). Overlay of hTRPV1_CAP_ and hTRPV1_6-Iodo-CAP_ structures (aligned with residues 416 to 712 from the transmembrane domain) shows that they bind in near-identical positions in the vanilloid pocket, with methoxy and hydroxyl vanilloid groups in the head region occupying the same positioning (Fig. 4f, g). 6-Iodo-CAP also makes extensive hydrophobic contacts similar to hTRPV1_CAP_ in the vanilloid pocket that include L515, L518, L547, L553, A566, I573, F587, F591, I662, L663, A666, and L670 (Fig. 3c, 4f). Consistent with the established TRPV1 vanilloid binding model, 6-Iodo-CAP and capsaicin both make hydrogen bonds with Y511 (Fig. 3c, 4f). Y511 is highly conserved among vertebrate TRPV1 orthologs and established as an important determinant of vanilloid recognition, further supporting the functional importance of this conserved interaction^14, 35, 37, 66^.

The most dramatic structural differences between hTRPV1_6-Iodo-CAP_ and hTRPV1_CAP_ occur at the S4-S5 linker, where 6-Iodo-CAP displaces the linker towards the pore. To place this displacement in the context of channel gating, we compared hTRPV1_6-Iodo-CAP_ to the experimentally determined open-state structure of rat TRPV1 (rTRPV1_Open_; PDB 5IRX), as hTRPV1_CAP_ and hTRPV1_6-Iodo-CAP_ both adopt closed conformations^32^. In this comparison, the 6-Iodo-CAP 6ʹ-iodide faces the S4-S5 linker and pushes the linker and S6 helix towards the pore by ∼2.0 Å and ∼1.4 Å, respectively (Fig. 4g). When compared to another open rat TRPV1 structure (PDB 7RQW) the same movements and distances are observed and the RMSD value is ∼1.00 Å for both open structures when the transmembrane domain of hTRPV1_6-Iodo-CAP_ is overlayed with the same region in the both open structures (Supplementary Fig. 3f). A structural morph between rTRPV1_Open_ (PDB 5IRX) and hTRPV1_6-Iodo-CAP_ highlights this displacement (Supplemental Movie 1). Previous cryo-EM studies and molecular dynamics simulations show that agonists, such as CAP, activate TRPV1 through pulling the S4–S5 linker away from the pore domain and channel gate^30, 32, 33, 58^. The opposite movement of the S4-S5 linker towards the S6 helix channel gate is observed in other TRPV1 antagonist-bound structures, including the allosterically modulating antagonist α-humulene and orthosteric antagonist SB-366792, and suggests a general mechanism for channel antagonism^28, 29, 32, 57^. This is further supported by a study that showed smaller halogens in this same position, result in a weaker antagonist, likely because the S4-S5 linker and S6 helix are moved to a lesser degree by a smaller substituent. hTRPV1_6-Iodo-CAP_ reveals that the iodide group sterically pushes I569 on the S4-S5 linker, which results in E570 movement away from R557 and breaks a previously proposed salt bridge required for channel activation (Fig. 4g,h, Supplementary Fig. 4g).

Moreover, R557 is free to hydrogen bond with the 6-Iodo-CAP hydroxyl group, which is not observed in agonist-bound structures (Fig. 4f, Supplementary Fig. 4g)^30, 33, 56, 58^. The requirement of breaking this salt bridge for inhibition is reported in structures of human TRPV1 bound to SB-366791 and SAF312 and functionally characterized by a double charge swap mutation (R557E/E570R) that partially rescued severe gating defects of a single mutation at either residue^28, 29, 67^. Furthermore, previously published NMR and electrophysiology experiments on the VSLD and full length channel indicate R557 is crucial for coupling temperature sensing to opening at the pore domain^16^.

Since 6-Iodo-CAP is the simplest conversion from an agonist to an antagonist, the structural observations show the minimum requirements for orthosteric antagonism which include interaction with Y511, disruption of the R557–E570 salt bridge and S4-S5 linker and S6 movement towards the pore (Supplementary Movie 1). These align with the inhibition requirements proposed for SB-366791-bound hTRPV1 by Neuberger et. al^29^. We further examine these requirements for antagonism through structural and functional analysis of TRPV1 antagonists with diverse chemical structures.

### Binding pocket plasticity enables diverse antagonist chemotypes

We performed a chemoinformatic multidimensional scaling (MDS) analysis to evaluate the chemical space that TRPV1 modulators occupy, and as expected, 6-Iodo-CAP is closest to capsaicin and dihydrocapsaicin in the MDS mathematical latent space (Fig. 1b, Supplementary Table 1). To further understand the mechanisms that diverse TRPV1 antagonist chemotypes use, we chose clinically relevant compounds Asivatrep, Mavatrep, and JNJ-17203212 based on their diversity as measured in MDS space and their distal relationship to other structurally characterized vanilloid binding site antagonists^28, 29^. Qualitatively, the most significant ligand structural differences appear in the so-called head and neck regions. Although these antagonists share a trifluoromethyl-containing tail, their neck and head groups differ substantially, including distinct six-membered ring systems and hydrogen bonding networks (Fig. 1c). We note that these antagonists have similar molecular volumes, where the overall size differences are relatively modest, ranging from less than one methyl group equivalent to roughly one and a half methyl groups. This diversity provides an opportunity to assess how the vanilloid pocket accommodates divergent chemical scaffolds of similar size while maintaining antagonism.

To define their binding modes, we determined hTRPV1 cryo-EM structures bound to Asivatrep (hTRPV1_Asivatrep_), Mavatrep (hTRPV1_Mavatrep_), and JNJ-17203212 (hTRPV1_JNJ-17203212_) at 2.1 Å, 2.4 Å, and 2.3 Å, respectively (Fig. 5a-d, Supplementary Fig. 5a, b, Supplementary Table 2). The high-resolution ligand densities allowed us to unambiguously model ligands in the vanilloid pocket and yield ligand Q-scores of 0.851, 0.806 and 0.835 for Asivatrep, Mavatrep, and JNJ-17203212, respectively (Supplementary Fig. 5c-e).

**Fig. 5:**
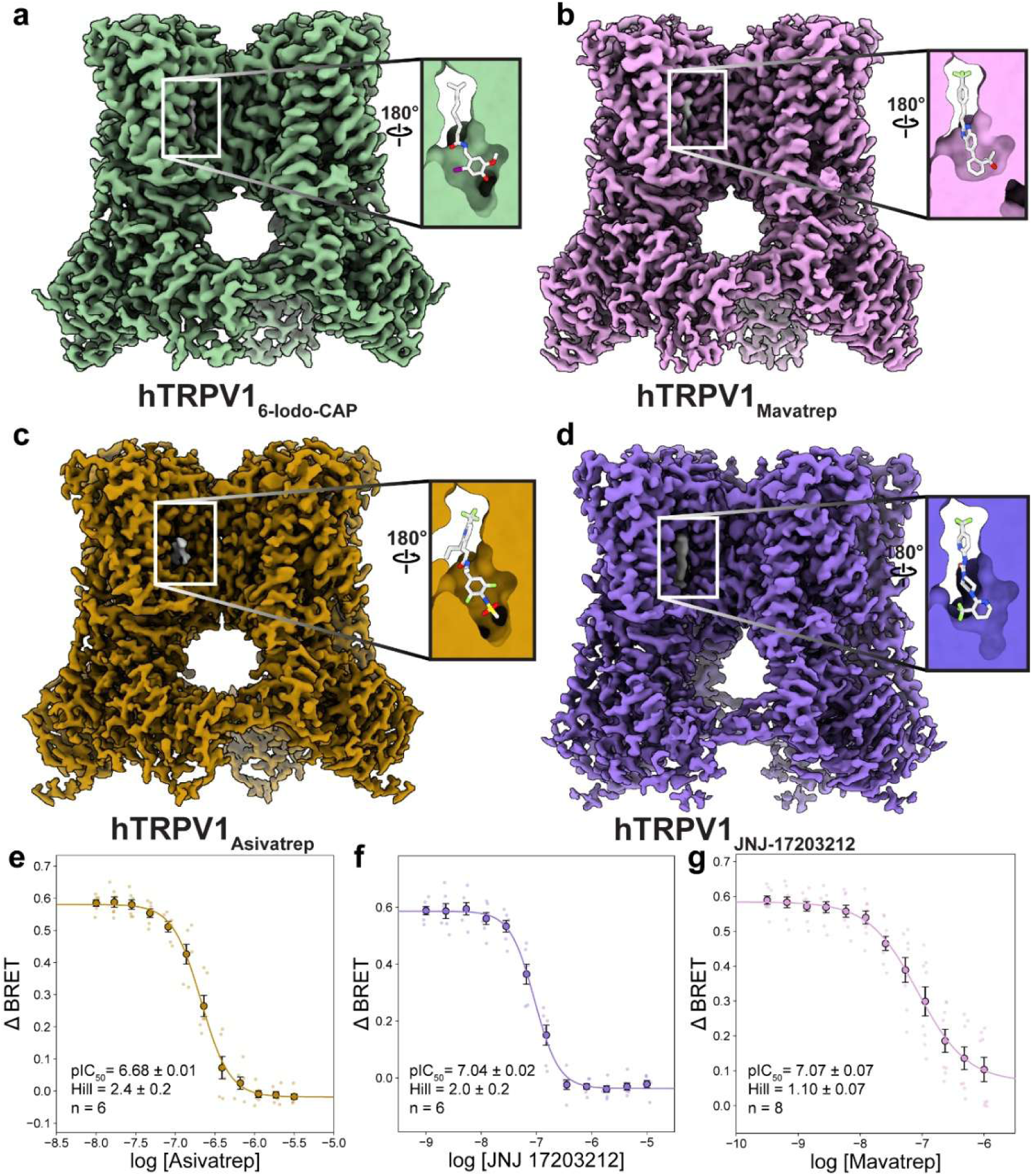
Ligand-dependent plasticity of the TRPV1 vanilloid pocket. Cryo-EM density for hTRPV16-Iodo-CAP (**a**), hTRPV1_Mavatrep_ (**b**), hTRPV1_Asivatrep_ (**c**), and hTRPV1_JNJ-17203212_ (**d**) and the inset shows the model represented as a surface viewed from inside-out the vanilloid pocket. The ligand model is displayed in white stick representation. BRET Ca^2+^ influx dose-response curves for Asivatrep (**e**), JNJ-17203212 (**f**), and Mavatrep (**g**) in the presence of 250 nM capsaicin show that, despite stabilizing distinct binding profiles, all efficiently inhibit TRPV1. Jittered data represent distinct replicates with the dose average values shown in a colored circle; the error bar is SEM. The p*IC*_50_ and n_H_ error values are derived from fitting errors.

The most striking difference in TRPV1 among these structures is the degree of vanilloid pocket plasticity. The top of the pocket where the tail group binds is relatively unchanged, with the width ∼13-14 Å across all structures; however, the width at the bottom of the pocket changes dramatically between ligands. While hTRPV1_6-Iodo-CAP_, hTRPV1_Asivatrep_, and hTRPV1_JNJ-17203212_ display an open cavity under the head group, hTRPV1_Mavatrep_ lacks this cavity. This vanilloid pocket plasticity observed in these structures and previously published SB-366791 and Libvatrep (SAF312)-bound structures still allows for pore closure by strikingly diverse chemotypes (Supplementary Fig. 6)^28, 29^. In hTRPV1_JNJ-17203212_, JNJ-17203212 is more exposed to the hydrophobic interior of the membrane compared to other ligand-bound TRPV1 structures, which suggests it may have a smaller interface with TRPV1 (Fig. 5d).

Despite the large deviations in the vanilloid pocket, all structures adopt a closed conformation with minimal variation in the pore (Supplementary Fig. 7). The primary deviation is localized to the intracellular end of the pore at residue N688, where the pore radius in hTRPV1_Asivatrep_ is ∼0.5 Å wider than hTRPV1_Mavatrep_ and hTRPV1_6-Iodo-CAP_, and ∼1 Å narrower in hTRPV1_JNJ-17203212_ (Supplementary Fig. 7). These findings indicate that structurally distinct antagonists converge on a common closed conformation.

To compare channel inhibition among the antagonists of interest under nearly identical conditions, BRET experiments were performed to measure inhibition of 250 nM CAP stimulated Ca^2+^ flux by Asivatrep, Mavatrep, and JNJ-17203212 (Fig. 5e-g). Because human TRPV1 exhibits baseline temperature activation at 37 °C, and distinct antagonist chemotypes are known to differentially inhibit this heat-evoked gating, evaluating these compounds at core physiological temperatures inherently conflates orthosteric ligand antagonism with allosteric heat inhibition^14^. Conducting the assays at 25 °C isolates the pure ligand-inhibition mechanism. When these experiments were repeated at 37 °C, compound-specific shifts in potency were observed (Supplementary Fig. 8). This variability indicates that these diverse scaffolds modulate the heat-activation pathway to varying degrees, supporting the 25°C assay strategy as an effective way to compare vanilloid-pocket target engagement accurately, independent of contributions from allosteric heat activation. All antagonists display high potency despite their diverse chemotypes, with Mavatrep p*IC*_50_ = 7.07 ± 0.07 (*IC*_50_ ≈ 85 nM), while Asivatrep p*IC*_50_ =6.68 ± 0.01, (*IC*_50_ ≈ 210 nM) and JNJ-17203212 p*IC*_50_ = 7.04 ± 0.02, (*IC*_50_ ≈ 90 nM) as shown in Fig. 5e-g. Despite substantial differences in vanilloid pocket architecture, these antagonists produce broadly similar functional inhibition. Mavatrep occupies a slightly more compact pocket with minimal cavity under the head group; whereas, Asivatrep and JNJ-17203212 leave a larger cavity in that region (Fig. 5a-d). Thus, the vanilloid pocket can accommodate chemically and structurally diverse antagonists while maintaining a similar functional outcome, although differences in pocket architecture may contribute to the modest differences in inhibitory potency observed among these compounds. All antagonists stabilize a closed pore, and these structural observations provide a framework to interpret specific molecular interactions of each ligand.

### Vanilloid pocket plasticity accommodates simultaneous ligand and lipid binding

The largest ligand-based deviation observed in ligand binding position is in hTRPV1_Asivatrep_ where the Asivatrep tail juts out of the top of the vanilloid pocket (Fig. 5c, 6a). In addition, the head group overlays well with other antagonists and the Asivatrep head group makes additional hydrogen bonds with the lower portion of the binding pocket (Fig. 6b). The two oxygens on the methylsulfonylamino group form hydrogen bonds with S512 and R557 and forms a hydrogen bond network between R557 and a water molecule (Fig. 6b, Supplemental Fig. 9). Another water molecule is observed to interact with E570 but not coordinated through Asivatrep (Fig. 6b). A third water molecule is 2.8 Å from T550 and 2.9 Å from the Asivatrep amide in the neck region (Supplementary Fig. 9c). In published structures, T550 is shown to be important for ligand binding, but previous structural studies have not shown a water-mediated interaction with ligand. T550 is 4.6 Å away from any hydrogen bonding groups in Asivatrep, a distance that does not correspond to a hydrogen bond. These water molecules are not observed in our other structures; however, it is possible that they are present in the hTRPV1_Asivatrep_ cryo-EM density map due to higher resolution reconstruction (2.10 Å) compared to other structures, which are at lower resolutions. Finally, the head group does not facilitate the interaction between R557 and E570 as they are 5.8 Å apart and these residues do not form a salt bridge (Fig. 6b). Apart from this Asivatrep bound structure, the R557–E570 salt bridge has been observed canonically in all TRPV1 antagonist-bound structures, and has been hypothesized to be required for channel _antagonism28, 29, 32, 57._

**Fig. 6:**
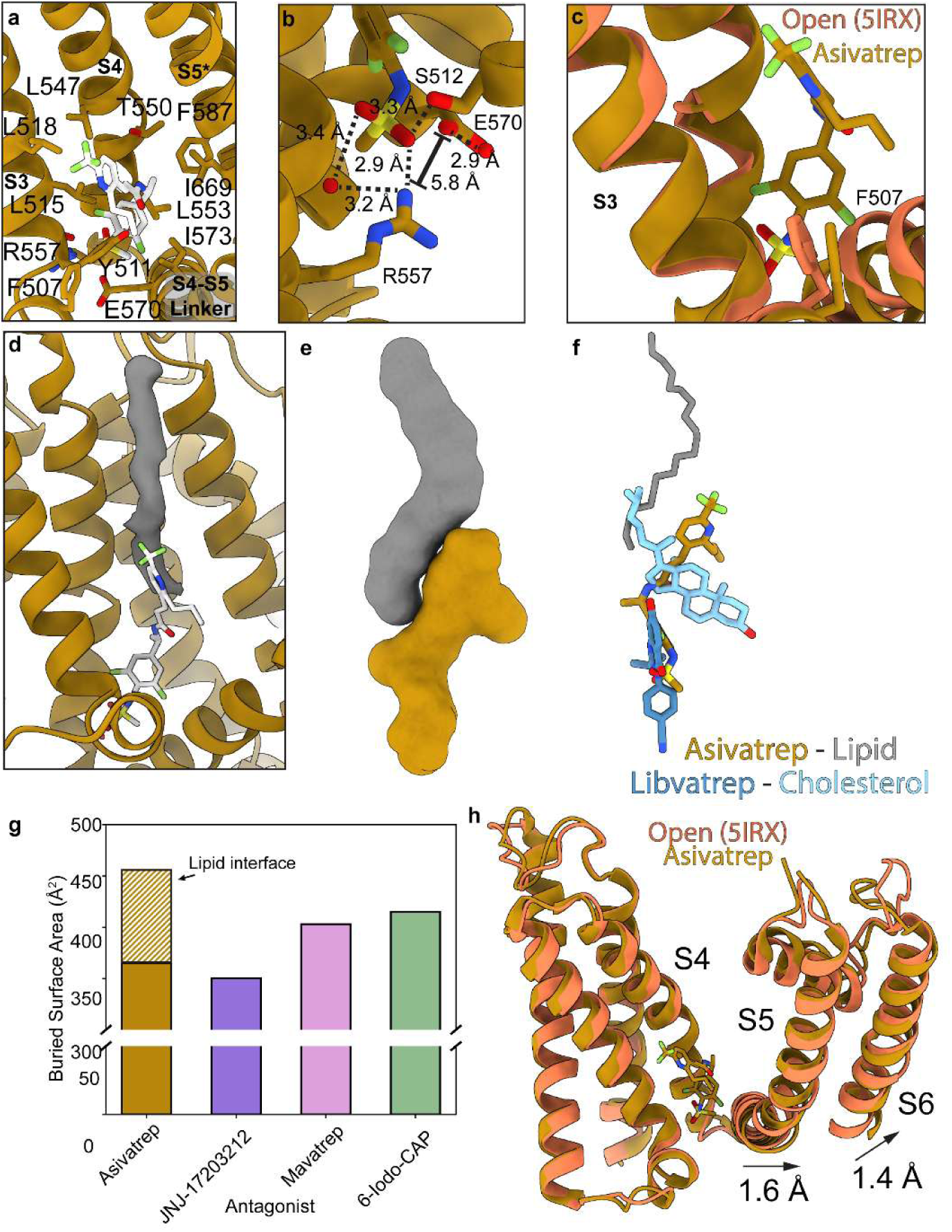
Simultaneous Asivatrep and lipid binding in the TRPV1 vanilloid pocket. **a** Vanilloid binding pocket for hTRPV1_Asivatrep_ with Asivatrep and interacting residues labeled and represented as sticks. **b** Zoomed in view at the bottom of the vanilloid pocket with Asivatrep interacting residues labeled and water molecules represented as red dots. Measured distances are displayed next to dashed lines (within 3.5 Å) or brackets (>3.5 Å). **c** Overlay between hTRPV1_Asivatrep_ (orange) and rTRPV1_Open_ (coral; PDB 5IRX) and F507 represented by sticks for both structures. Alignment was performed between the transmembrane domains (residues 416-712). **d** Difference map (grey) between hTRPV1_Asivatrep_ cryo-EM density and model generated map. **e** Asivatrep (orange) and eicosane (grey) model surface representation. **f** Ligand overlay for hTRPV1_Asivatrep_ (Asivatrep orange, eicosane grey) and hTRPV1_Libvatrep_ (Libvatrep dark blue, cholesterol light blue) where all structures are aligned via the transmembrane domain. **g** Buried surface area calculated in ChimeraX of TRPV1 antagonists in the corresponding structures and Asivatrep-lipid only interface (orange diagonal lines) **h** Overlay of hTRPV1_Asivatrep_ (orange) and rTRPV1_CAP_ (coral; PDB 5IRX) with arrows indicating hTRPV1_Asivatrep_ movement relative to rTRPV1_Open_. Alignment performed same as (**c**).

In addition to hydrogen bonds, there are hydrophobic interactions that primarily take place in the tail group and they include F507, L515, L518, L547, L553, I573, F587, and L670 (Fig. 6a). A hydrophobic interaction unique to hTRPV1_Asivatrep_, is between F507 on the linker between S2 and S3 and the propyl group that is attached to the Asivatrep tail (Fig. 6a, c). This interaction is not observed in other ligand-bound TRPV1 structures and an overlay between rTRPV1_Open_ (5IRX) and hTRPV1_Asivatrep_ aligned by the transmembrane domain shows F507 movement towards the Asivatrep to accommodate this interaction (Fig. 6c).

Lastly, an unidentified tube-like density occupies the space in the vanilloid pocket where Asivatrep does not interact. This density looks similar to lipid acyl-chain density and is clearly not endogenous cholesterol (Fig. 6d). Our apo structure in GDN and other TRPV1 structures solubilized in detergents confirms endogenous lipids bind tightly enough to hTRPV1 to be copurified even in a detergent micelle. Since our hTRPV1_Asivatrep_ dataset is relatively large (∼40K micrographs), we attempted extensive 3D classification and 3D variability analysis to determine if the lipid density observed in hTRPV1_Asivatrep_ is from the same lipid that occupies the vanilloid pocket in apo structures.

However, this analysis showed the primary structural variability in our dataset comes from the noise in the micelle and the ankyrin flexibility, and the reported structure is the predominant conformation in the dataset. While we cannot identify the molecule in this density, the hTRPV1 structure bound to Libvatrep shows similarity to hTRPV1_Asivatrep_ where Libvatrep sits in the lower region of the vanilloid pocket and a cholesterol co-binds at the site where we observe this potential lipid density (Fig. 6f, Supplementary Fig. 6a)^28^. It may be possible in future studies to determine the identity of this co-bound lipid molecule.

A full lipid molecule could not be modeled into this density, but to understand how an annular lipid contributes to Asivatrep binding a 20-carbon alkane (eicosane) was modeled into the density (Q_Ligand_ = 0.822). The contact between the modeled alkane and Asivatrep is 90 Å^2^, whereas the contact between hTRPV1 and Asivatrep is 416 Å^2^ (Fig. 6e, g). The combined interactions between the alkane and hTRPV1 to Asivatrep is 506 Å^2^ which is greater than the surface area contacts between hTRPV1_6-iodo-CAP_ (472 Å^2^), hTRPV1_mavatrep_ (468 Å^2^), and hTRPV1_JNJ-17203212_ (414 Å^2^) (Fig. 6g). With increased contact with an annular lipid, it is likely Asivatrep has a higher affinity for hTRPV1 than 6-iodo-dihydrocapsaicin, Mavatrep or JNJ-17203212 despite the tail group sticking out of the binding pocket.

Although the Asivatrep tail group does not occupy the top of the vanilloid pocket, its binding still induces a shift of the S4–S5 linker and S6 helix towards the pore, consistent with our other structures and previously reported antagonist-bound TRPV1 conformations such as hTRPV1_SB-366791_ and hTRPV1_SAF312_ (Fig. 6h, Supplementary Movie 2)^28, 29^. Notably, this occurs despite differences in ligand engagement within the pocket, which suggests that the engagement of the ligand tail within the upper pocket is not strictly required for these conformational changes, and that interactions at the base of the pocket are sufficient to stabilize the closed state.

### hTRPV1 antagonism does not require R557 and E570 salt bridge disruption

The vanilloid pocket in hTRPV1_Mavatrep_ exhibits substantial contraction compared to other ligand-bound structures and lacks the lower cavity that is typically observed beneath antagonist-bound structures (Fig. 5b). Within this compact pocket, Mavatrep engages in conserved interactions that include a hydrogen bond between its neck amide and Y511, consistent with all reported orthosteric ligand-bound hTRPV1 structures (Fig. 7a, Supplementary Fig. 10)^28, 29, 30, 32, 33, 56, 58^. Additionally, T550 makes an indirect interaction with Mavatrep via a hydrogen bond with a water molecule similar to hTRPV1_Asivatrep_ (Fig. 7b, Supplementary Fig. 10b). The ligand is further stabilized by extensive hydrophobic contacts that involve residues L515, F543, A546, L547, L553, A566, I569, V583, F587, F591, I662, L663, A666, and L670 (Fig. 7a). Like Asivatrep and 6-Iodo-CAP, Mavatrep also hydrogen bonds with R557, which is proposed to disrupt the R557–E570 salt bridge interaction in the SAF-312-bound hTRPV1 structure (Fig. 7c)^28^. However, the interaction between Mavatrep and R557 stabilizes the R557 side chain in a position closer to the S4–S5 linker and is further stabilized by a hydrogen bond with S512 which is 3.5 Å away from R557 (Fig. 7c). In this position enforced by Mavatrep, R557 makes an electrostatic interaction with E570, yet it inhibits hTRPV1 (Fig. 5g, 7c). This is in stark contrast to previous cryo-EM structures where the R557–E570 salt bridge is intact in open structures and absent in apo and antagonist-bound structures^28, 29, 30, 32, 33, 34, 56, 57, 58^. These observations demonstrate that channel closure is not strictly dependent on the disruption of the R557–E570 interaction despite what is seen with 6-Iodo-CAP, Asivatrep, SB-366791, and SAF312 binding^28, 29^. Structural comparison with rTRPV1_Open_ overlayed by the transmembrane domain (residues 416-712) shows close alignment of the voltage sensing-like domain (VSLD; S1–S4; RMSD 2.2 Å), with movement localized at the bottom of S3 and the S4–S5 linker (2.2 Å) (Fig. 7d, Supplementary Movie 3). In contrast to other antagonist-bound structures, which show minimal rearrangement in S3, these changes for hTRPV1_Mavatrep_ likely arise from a distinct interaction network that involves Mavatrep, S512, R557, and E570 that couples movement of the bottom of S3 to displacement of the S4–S5 linker and stabilizes the closed pore.

**Fig. 7:**
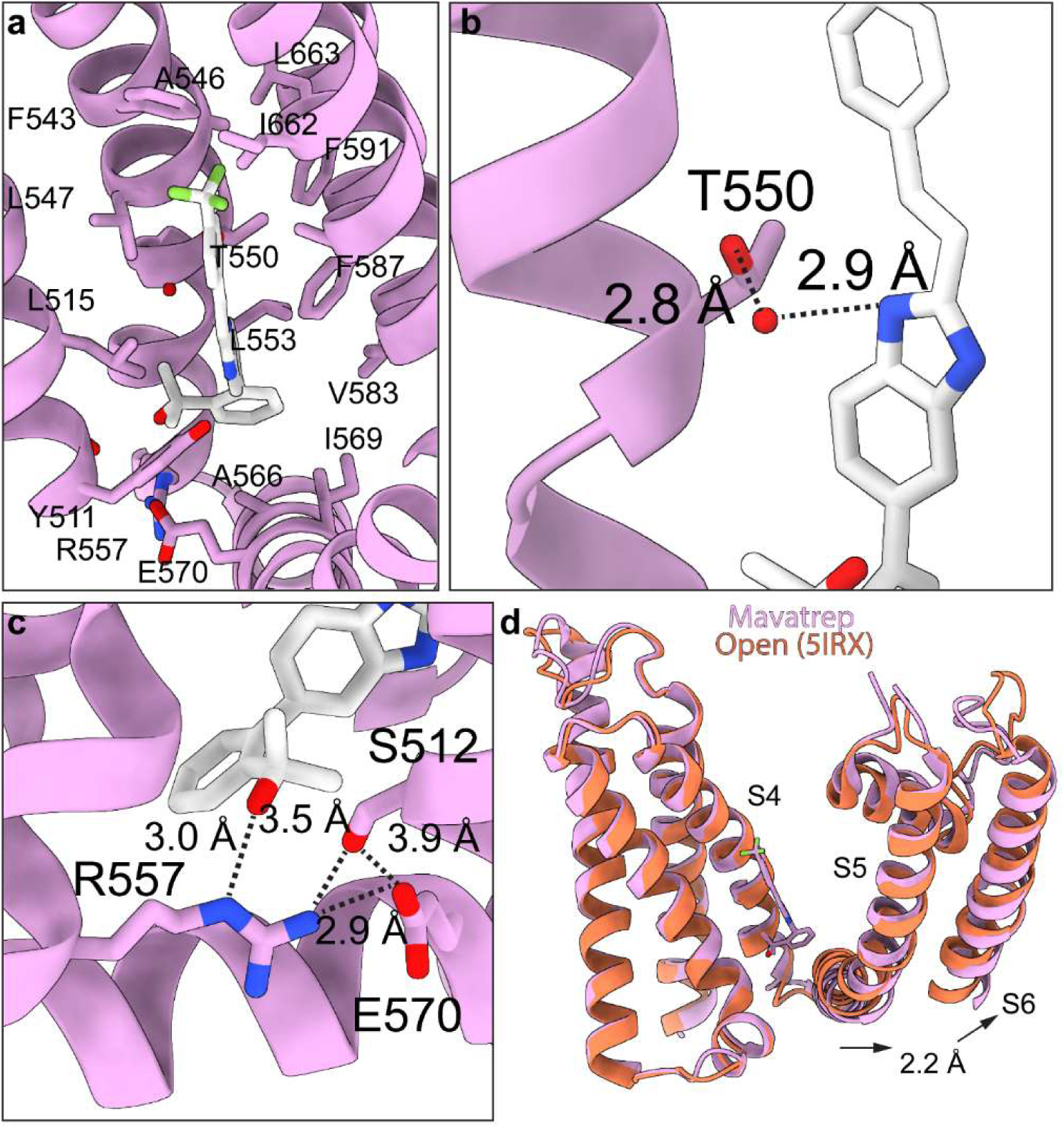
Mavatrep maintains the R557-E570 interaction in hTRPV1. **a** Vanilloid binding pocket for hTRPV1_Mavatrep_ with Mavatrep and interacting residues labeled and represented as sticks. **b** Zoomed in view of T550 and water (red dot) interaction with Mavatrep neck group. Cryo-EM density for water molecule shown with mesh and distances with dashed lines. **c** Zoomed in view of R557 and E570 salt bridge interaction in hTRPV1_Mavatrep_ with corresponding interactions labeled and displayed by dashed lines. **d** Overlay of hTRPV1_Mavatrep_ (pink) and rTRPV1_Open_ (coral; PDB 5IRX) with corresponding movements shown with arrows. Alignment was performed between the transmembrane domains (residues 416-712).The vanilloid pocket in hTRPV1_Mavatrep_ exhibits substantial contraction compared to other ligand-bound structures and lacks the lower cavity that is typically observed beneath antagonist-bound structures (Fig. 5b). Within this compact pocket, Mavatrep engages in conserved interactions that include a hydrogen bond between its neck amide and Y511, consistent with all reported orthosteric ligand-bound hTRPV1 structures (Fig. 7a, Supplementary Fig. 10)^28, 29, 30, 32, 33, 56, 58^. Additionally, T550 makes an indirect interaction with Mavatrep via a hydrogen bond with a water molecule similar to hTRPV1_Asivatrep_ (Fig. 7b, Supplementary Fig. 10b). The ligand is further stabilized by extensive hydrophobic contacts that involve residues L515, F543, A546, L547, L553, A566, I569, V583, F587, F591, I662, L663, A666, and L670 (Fig. 7a). Like Asivatrep and 6-Iodo-CAP, Mavatrep also hydrogen bonds with R557, which is proposed to disrupt the R557–E570 salt bridge interaction in the SAF-312-bound hTRPV1 structure (Fig. 7c)^28^. However, the interaction between Mavatrep and R557 stabilizes the R557 side chain in a position closer to the S4–S5 linker and is further stabilized by a hydrogen bond with S512 which is 3.5 Å away from R557 (Fig. 7c). In this position enforced by Mavatrep, R557 makes an electrostatic interaction with E570, yet it inhibits hTRPV1 (Fig. 5g, 7c). This is in stark contrast to previous cryo-EM structures where the R557–E570 salt bridge is intact in open structures and absent in apo and antagonist-bound structures^28, 29, 30, 32, 33, 34, 56, 57, 58^. These observations demonstrate that channel closure is not strictly dependent on the disruption of the R557–E570 interaction despite what is seen with 6-Iodo-CAP, Asivatrep, SB-366791, and SAF312 binding^28, 29^. Structural comparison with rTRPV1_Open_ overlayed by the transmembrane domain (residues 416-712) shows close alignment of the voltage sensing-like domain (VSLD; S1–S4; RMSD 2.2 Å), with movement localized at the bottom of S3 and the S4–S5 linker (2.2 Å) (Fig. 7d, Supplementary Movie 3). In contrast to other antagonist-bound structures, which show minimal rearrangement in S3, these changes for hTRPV1_Mavatrep_ likely arise from a distinct interaction network that involves Mavatrep, S512, R557, and E570 that couples movement of the bottom of S3 to displacement of the S4–S5 linker and stabilizes the closed pore.

### JNJ-17203212 antagonizes TRPV1 without Y511 interaction

As opposed to most antagonists, JNJ-17203212 makes only a single hydrogen bond with T550 and is engaged primarily in hydrophobic interactions with L515, F543, A546, L547, L553, I569, I573, F587, F591, I662, L663, and L670 (Fig. 8a, Supplementary Fig. 11). Despite sparce hydrophobic interactions, the *IC*_50_ for JNJ-17203212 is 91 nM, comparable to Asivatrep (209 nM) and more potent than 6-Iodo-CAP (427 nM) (Fig. 4c, 5e-g).

**Fig. 8:**
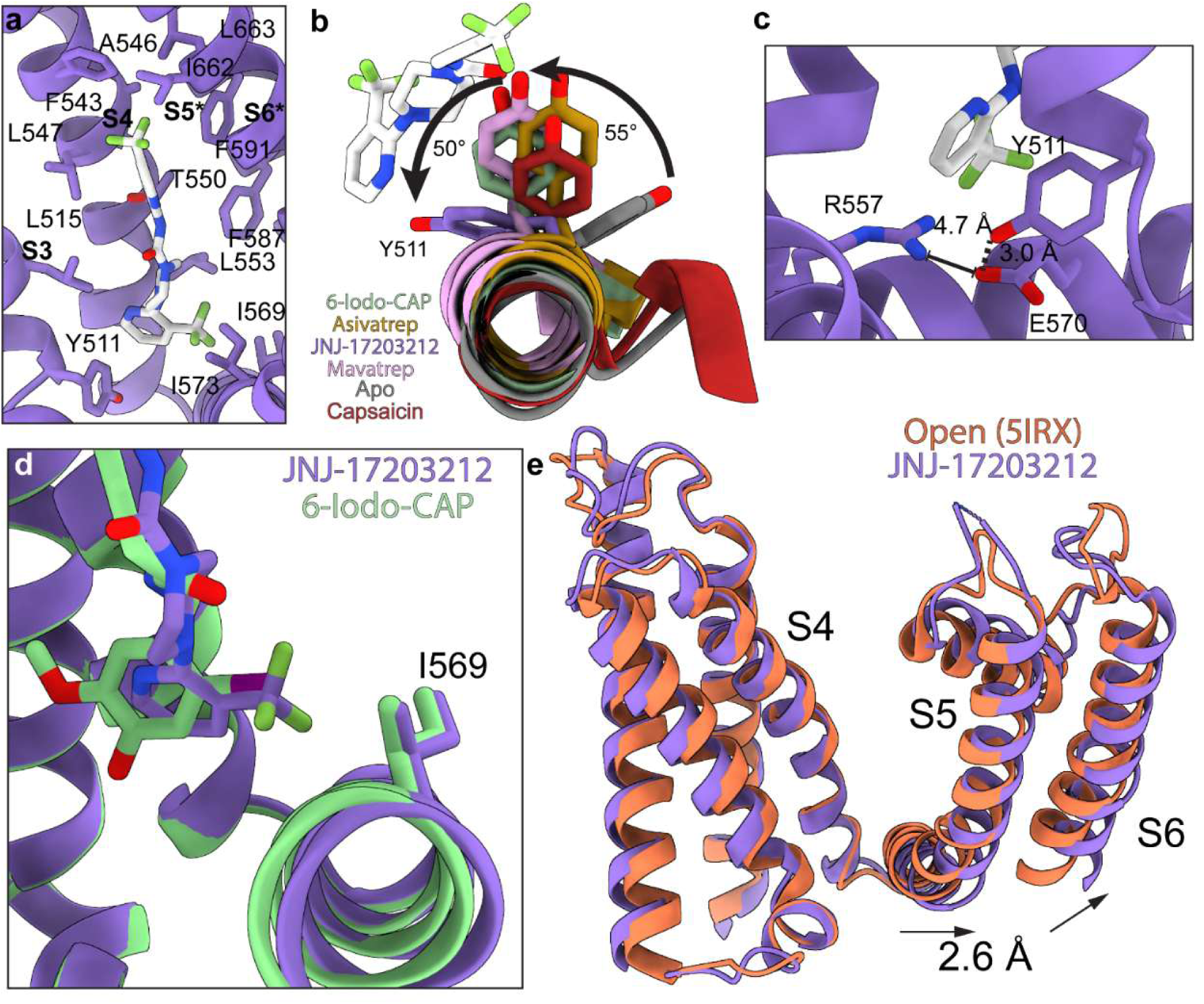
JNJ-17203212 does not interact with Y511 in hTRPV1. **a** Vanilloid binding pocket for hTRPV1_JNJ-17203212_ with JNJ-17203212 and interacting or important residues represented as sticks. **b** hTRPV1_6-Iodo-CAP_ (green), hTRPV1_Asivatrep_ (orange), hTRPV1_JNJ-17203212_ (purple), hTRPV1_Mavatrep_ (pink), hTRPV1_Apo_ (grey), and sqTRPV1_Capsaicin_ (red) aligned by the transmembrane domain and JNJ-17203212 displayed as stick representation viewed down the S3 helix and Y511 shown as sticks for all structures. Relative rotations displayed as a arrows and labeled with the degrees of rotation. **c** Zoomed in view of the bottom of the vanilloid pocket with R557 and E570 displayed as sticks and their distance measured and indicated with dashed lines. **d** Overlay between hTRPV1_6-Iodo-CAP_ (green) and hTRPV1_JNJ-17203212_ (purple) aligned via the transmembrane domain (residues 416-712) and the S4-S5 linker (ribbon) and I569 (sticks) are displayed for both structures. **e** Overlay of hTRPV1_JNJ-17203212_ (purple) and sqTRPV1_Capsaicin_ (grey) with corresponding movements shown with arrows. Alignment performed same as (**d**).

Unexpectedly, JNJ-17203212 does not engage Y511, a residue reported to contribute to antagonist binding in hTRPV1, specifically SB-366791 and SAF312 (Fig. 8a)^28, 29^. The Y511 hydroxyl oxygen is 4.0 Å away from the JNJ-17203212 head group amide, a distance inconsistent with hydrogen bonding (Fig. 8b). Consistent with the lack of interaction, the Y511 side chain is rotated ∼50° from other ligand-bound structures and ∼105° rotated relative to Y511 in hTRPV1_Apo_(Fig 8b, Supplementary Movie 4). The altered Y511 conformation highlights the vanilloid pocket plasticity and suggests Y511 can accommodate ligands without direct participation in ligand recognition. Although the JNJ-17203212 conjugated head group could potentially support a π stacking interaction, the cryo-EM density places Y511 in a geometry incompatible with a π-π interaction. Instead, Y511 forms a hydrogen bond with E570, an interaction not observed in other ligand-bound structures (Fig. 8c, Supplementary Fig. 11b).

Despite the absence of Y511 engagement, JNJ-17203212 stabilizes the closed channel conformation similar to other antagonist-bound structures (Supplementary Fig. 7). The trifluoromethyl on the head group extends towards the S4-S5 linker and displaces I569 in a manner reminiscent of the iodo moiety in 6-Iodo-CAP (Fig 8d). This interaction shifts I569 and the S4-S5 linker ∼2.3 Å toward the pore relative to hTRPV1_Open_, and the S6 helix also moves ∼2.3 Å inward. This movement represents the largest pore-directed displacement among the antagonists characterized here (Fig. 8e, Supplemental Movie 4). These observations indicate that Y511 engagement is not a strict requirement for TRPV1 antagonism.

### Targeted mutagenesis validates ligand-specific interactions

The structures presented in this study reveal distinct ligand-specific interaction networks within the orthosteric vanilloid binding pocket. To validate that the variable interactions identified from our cryo-EM structural studies are functionally meaningful, we evaluated mutated residues that should have variable functional effect that depends on the particular antagonist. We selected Y511F, T550V, F507A, and I569A based on their cryo-EM identified interactions with the antagonists, and, in the case of F507A and I569A, the identification of previously uncharacterized interactions within the vanilloid binding pocket. Because these mutations may also alter the intrinsic activation properties of hTRPV1, we first characterized the BRET response of each mutant to capsaicin before antagonist activity assessment.

Capsaicin dose-response curves were recorded for Y511F, T550V, F507A, and I569A to determine whether each mutant retained capsaicin-dependent channel activation (Fig. 9a). Only three mutants remain responsive to capsaicin, although the maximal response is reduced relative to WT. Y511F, T550V, and F507A reached approximately 50%, 80%, and 100%, of the maximal WT response, respectively (Supplementary Fig. 12a). The mutant I569A was not responsive to capsaicin concentrations up to 10 µM (Data not shown), which is consistent with previously reported results^68^. In addition to changes in maximal response, capsaicin potency is reduced for each mutant with pEC_50_ values of 8.34 ± 0.06 for WT, 7.69 ± 0.12 for Y511F, 7.13 ± 0.03 for T550V, and 8.56 ± 0.03 for F507A (Table 1). Despite these changes in potency and maximal response, the Hill coefficients were largely unchanged between WT and the mutants, which indicates that the mutations do not substantially alter cooperativity (Fig. 9a, Table 1).

**Fig. 9:**
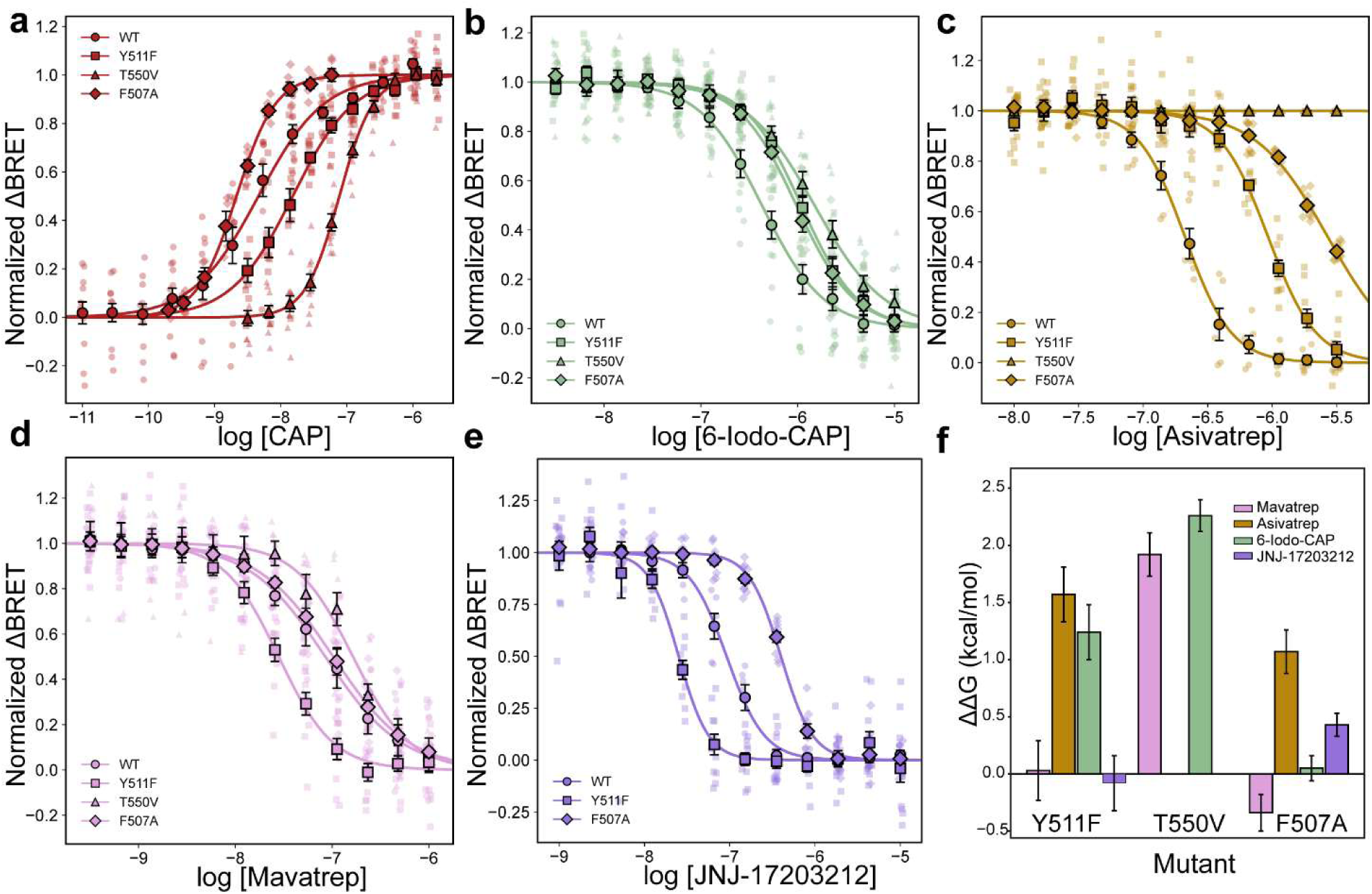
Targeted mutagenesis reveals ligand-specific effects on hTRPV1 antagonism. BRET dose response curves for capsaicin (**a**), 6-Iodo-CAP (**b**), Asivatrep (**c**), Mavatrep (**d**), and JNJ-17203212 (**e**) with wildtype (circles), Y511F (squares), T550V (triangles), and F507A (diamonds) datapoints shown. Jittered data correspond to the construct marker shape and represent distinct replicates with the dose average values shown in the solid, outlined marker. All antagonist experiments were performed with 250 nM capsaicin present. The error bar is SEM. Data were normalized independently for each construct to the minimum and maximum responses obtained from the corresponding four-parameter logistic fit, with the minimum response set to 0 and the maximum response set to 1. **f** Bar plot for ΔΔG values calculated from Cheng-Prusoff analysis for each antagonist and all three mutants. Error was calculated as described in the methods.

**Table 1.** Pharmacological and thermodynamic parameters of TRPV1 antagonists across wild-type and binding-site mutant constructs.

| Ligand | Mutant | Replicates (n) | Hill Coefficient | pEC<sub>50</sub>/pIC<sub>50</sub> | Apparent K<sub>i</sub> (nM) | ΔΔG (kcal/mol) | Distance (Å)** |
| --- | --- | --- | --- | --- | --- | --- | --- |
| Capsaicin | WT | 12 | 0.87 ± 0.09 | 8.34 ± 0.06 | - | - | - |
|  | Y511F | 10 | 0.94 ± 0.23 | 7.82 ± 0.17 | - | - | 2.7 |
|  | T550V | 11 | 1.69 ± 0.17 | 7.11 ± 0.03 | - | - | 4.1 |
|  | F507A | 6 | 1.47 ± 0.17 | 8.67 ± 0.04 | - | - | - |
| 6-Iodo-CAP | WT | 12 | 1.36 ± 0.07 | 6.37 ± 0.02 | 7.66 ± 1.10 | 0 | - |
|  | Y511F | 11 | 1.49 ± 0.17 | 5.96 ± 0.04 | 62.6 ± 23.8 | 1.24 ± 0.24 | 2.7 |
|  | T550V | 10 | 1.20 ± 0.20 | 5.83 ± 0.08 | 351 ± 67.3 | 2.26 ± 0.14 | 4.2 |
|  | F507A | 10 | 1.45 ± 0.16 | 6.01 ± 0.04 | 8.29 ± 1.08 | 0.05 ± 0.11 | - |
| Asivatrep | WT | 6 | 2.4 ± 0.2 | 6.68 ± 0.01 | 3.75 ± 0.52 | 0 | - |
|  | Y511F | 11 | 2.33 ± 0.33 | 6.03 ± 0.03 | 53.3 ± 20.0 | 1.57 ± 0.24 | 2.9 |
|  | T550V* | 13 | - | - | - | - | via H<sub>2</sub>O |
|  | F507A | 10 | 1.56 ± 0.21 | 5.57 ± 0.12 | 22.8 ± 6.64 | 1.07 ± 0.19 | - |
| Mavatrep | WT | 6 | 1.10 ± 0.07 | 7.07 ± 0.07 | 1.53 ± 0.32 | 0 | - |
|  | Y511F | 11 | 1.53 ± 0.21 | 7.55 ± 0.04 | 1.61 ± 0.61 | 0.03 ± 0.26 | 3.4 |
|  | T550V | 8 | 1.47 ± 0.38 | 6.78 ± 0.10 | 39.3 ± 9.3 | 1.92 ± 0.19 | via H<sub>2</sub>O |
|  | F507A | 10 | 1.10 ± 0.19 | 6.99 ± 0.06 | 0.87 ± 0.14 | -0.34 ± 0.16 | - |
| JNJ-17203212 | WT | 8 | 2.0 ± 0.2 | 7.04 ± 0.02 | 1.64 ± 0.23 | 0 | - |
|  | Y511F | 11 | 2.48 ± 0.54 | 7.60 ± 0.04 | 1.43 ± 0.54 | -0.080 ± 0.24 | 4.0 |
|  | T550V* | 10 | - | - | - | - | 3.5 |
|  | F507A | 10 | 2.33 ± 0.19 | 6.40 ± 0.02 | 3.38 ± 0.34 | 0.43 ± 0.10 | - |

\*Constructs for which the concentration-response curve did not support a reliable midpoint estimate were excluded from the Cheng–Prusoff and Δ analyses
\*\*Distance between ligand and closest heteroatom on mutated residue measured from cryo-EM models. via H<sub>2</sub>O indicates a water-mediated interaction between hTRPV1 and ligand.

To quantitatively compare antagonist activity across WT and mutant hTRPV1, we calculate apparent binding affinity (*K_i_*) from antagonist IC_50_ measurements with a Cheng-Prusoff analysis^63, 64^. Because the mutations alter capsaicin potency, the capsaicin EC_50_ determined for each individual mutant was used in the corresponding calculation, together with the concentration of capsaicin used in the antagonist concentration-response experiment (see Methods). The change in apparent binding free energy (ΔΔG) was calculated from the change in *K_i_* values between the mutant and WT. This approach accounts for changes in capsaicin activation profiles caused by the mutations and provides a common energetic framework for evaluating the effects of each mutation on antagonist activity.

Mutation of Y511 to phenylalanine produced ligand-specific changes in antagonist apparent affinity. Y511F substantially increased the apparent *K_i_* values for 6-Iodo-CAP and Asivatrep, corresponding to ΔΔG values of 1.24 ± 0.24 kcal/mol and 1.57 ± 0.24 kcal/mol, respectively, consistent with a contribution of the Y511 hydrogen-bonding interaction to the activity of these antagonists (Fig. 9b, c, f, Table 1). In agreement with these functional measurements, Y511 forms a hydrogen bond with both 6-Iodo-CAP and Asivatrep and in their respective structures, with donor-acceptor distances of 3.2 Å and 2.9 Å between heteroatoms, respectively (Table 1). In contrast, Y511F has a negligible effect on Mavatrep (ΔΔG = 0.03 ± 0.26 kcal/mol) and JNJ-17203212 (ΔΔG = -0.08 ± 0.24 kcal/mol) (Fig. 9d, e, f, Table 1). Although Y511 appears to form a hydrogen bond with Mavatrep in the structure, the greater distance between the Y511 hydroxyl group and the corresponding hydrogen-bond acceptor (3.4 Å) may indicate a weaker interaction relative to 6-Iodo-CAP and Asivatrep (Table 1). Consistent with this minimal functional effect, Y511 is not positioned to form a direct interaction with JNJ-17203212 in the structure (Fig. 8a,b). Together, these results demonstrate that the functional consequence of Y511 mutation is strongly ligand dependent, with substantial energetic effects for 6-Iodo-CAP and Asivatrep, but no contributions for Mavatrep and JNJ-17203212 interactions.

Mutation of T550 to a valine results in a substantial reduction in capsaicin potency and strongly affects antagonist activity (Fig. 9a-e, Table 1). Reliable concentration-response fits were obtained for Mavatrep and 6-Iodo-CAP, with pIC_50_ values of 6.74 ± 0.05 and 5.85 ± 0.07 respectively, whereas the concentration-response curves for Asivatrep and JNJ-17203212 were shifted sufficiently so that reliable fits could not be determined (Fig. 9b-e, Supplementary Fig. 12, Table 1). For antagonists with reliable fits, Cheng-Prusoff analysis yields apparent *K_i_* for 6-Iodo-CAP of 351 ± 70 nM and for Mavatrep of 39.3 ± 9.3 nM, which corresponds to ΔΔG values of 2.26 ± 0.14 kcal/mol and 1.92 ± 0.19 kcal/mol, respectively (Fig. 9f, Table 1). These energetic effects are consistent with an important contribution of the T550 hydrogen bond network to antagonist activity.

The structural observations provide context for the different interactions that involve T550 across the antagonist-bound structures. T550 interacts with Asivatrep and Mavatrep through hydrogen bond networks that are mediated by a water molecule, whereas an ordered water molecule is not observed near T550 in hTRPV1_6-Iodo-CAP_, hTRPV1_JNJ-17203212_, or hTRPV1_CAP_ (Fig. 7b, Supplementary Fig. 9c).

Notably, the T550 hydroxyl group is outside conventional hydrogen bond distance from 6-Iodo-CAP (4.2 Å) and capsaicin (4.1 Å) despite the strong functional effect of T550V on activity of these two ligands.

The lower resolution of hTRPV1_6-Iodo-CAP_ (2.9 Å) and hTRPV1_CAP_ (3.1 Å) compared to hTRPV1_Asivatrep_ (2.1 Å) and hTRPV1_Mavatrep_ (2.4 Å) structures may limit the ability to resolve ordered water molecules in these complexes. Alternatively, the functional effect of T550V may reflect changes in local pocket interactions or conformational dynamics that are not directly resolved in the structures.

F507 represents a potentially novel ligand interaction identified in this study. F507 is positioned to form a hydrophobic interaction with Asivatrep and this interaction is not observed in any other published TRPV1 ligand-bound structures (Fig. 6c). Mutation of F507 to alanine has relatively little effect on capsaicin potency, while the the pIC_50_ values for 6-Iodo-CAP and Mavatrep were 6.00 ± 0.04 and 6.96 ± 0.07, respectively (Fig. 9a-c, Table 1). Cheng-Prusoff analysis yields ΔΔG values of 0.05 ± 0.11 kcal/mol and -0.34 ± 0.16 kcal/mol respectively (Fig. 9f, Table 1). The minimal energetic effect of F507A on 6-Iodo-CAP suggests F507 makes little contribution to its activity, whereas the negative ΔΔG for Mavatrep may indicate removal of the phenylalanine side chain slightly increases Mavatrep binding affinity. In contrast, F507A produces a modest reduction in the binding energy of JNJ-17203212, with a ΔΔG of 0.43 ± 0.10 kcal/mol (Fig. 9f, Table 1). Although F507 is not positioned to form a direct interaction with JNJ-17203212 in the structure, this effect raises the possibility that F507 influences local pocket dynamics or conformational states that affect JNJ-17203212 binding or activity. Indeed, previous studies highlight the importance of S2–S3 linker mobility in ligand and temperature activation^16, 69, 70^. Conversely, F507 side chain removal may favor conformations compatible with Mavatrep binding, consistent with the small decrease in apparent *K_i_* observed for Mavatrep. These effects highlight that mutations can influence antagonist activity through mechanisms beyond disruption of direct ligand-residue contact.

The most pronounced effect of F507A was observed for Asivatrep. The pIC_50_ decreased to 5.62 ± 0.08, which corresponds to an apparent *K_i_* of 22.8 ± 6.6 nM and a ΔΔG of 1.07 ± 0.19 kcal/mol (Fig. 9c, f, Table 1). This energetic penalty is consistent with a contribution of the F507–Asivatrep hydrophobic interaction to antagonist activity. Together, the functional effects of F507A support the structural identification of F507 as an interacting residue with Asivatrep, and this residue may influence the conformational properties of the pocket.

Together, these mutagenesis experiments demonstrate that mutated residues identified in our structures produce ligand-dependent functional effects on antagonism. Rather than acting as universal determinants of antagonist binding, individual residues produce distinct energetic effects that depend on the ligand and its interaction network within the vanilloid pocket. These functional data therefore provide experimental support for the ligand-specific interactions identified in our structures.

### Diverse antagonist interactions converge on a common inhibited state

Clinically relevant TRPV1 antagonists occupy a broad chemical diversity with distinct protein-ligand interaction networks (Fig. 1c,b, 5a-d), yet the distinct chemotypes are accommodated within the vanilloid pocket through binding site plasticity. Despite this chemical diversity, binding patterns, and binding site plasticity, antagonist-bound TRPV1 structures converge on a common inhibited conformation characterized by pore closure (Fig 5a-d, Supplemental Fig. 7). Notably, despite their diverse chemotypes and interaction networks, all compounds function as potent TRPV1 antagonists (Fig. 5e-g). Structural overlay of antagonist-bound complexes identifies that, not only does the vanilloid binding pocket rearrange to accommodate the distinct chemotypes, but also that the ligand poses exhibit substantial variability within the orthosteric vanilloid pocket, with pronounced differences in both head and tail group occupancy (Fig. 10a).

**Fig. 10:**
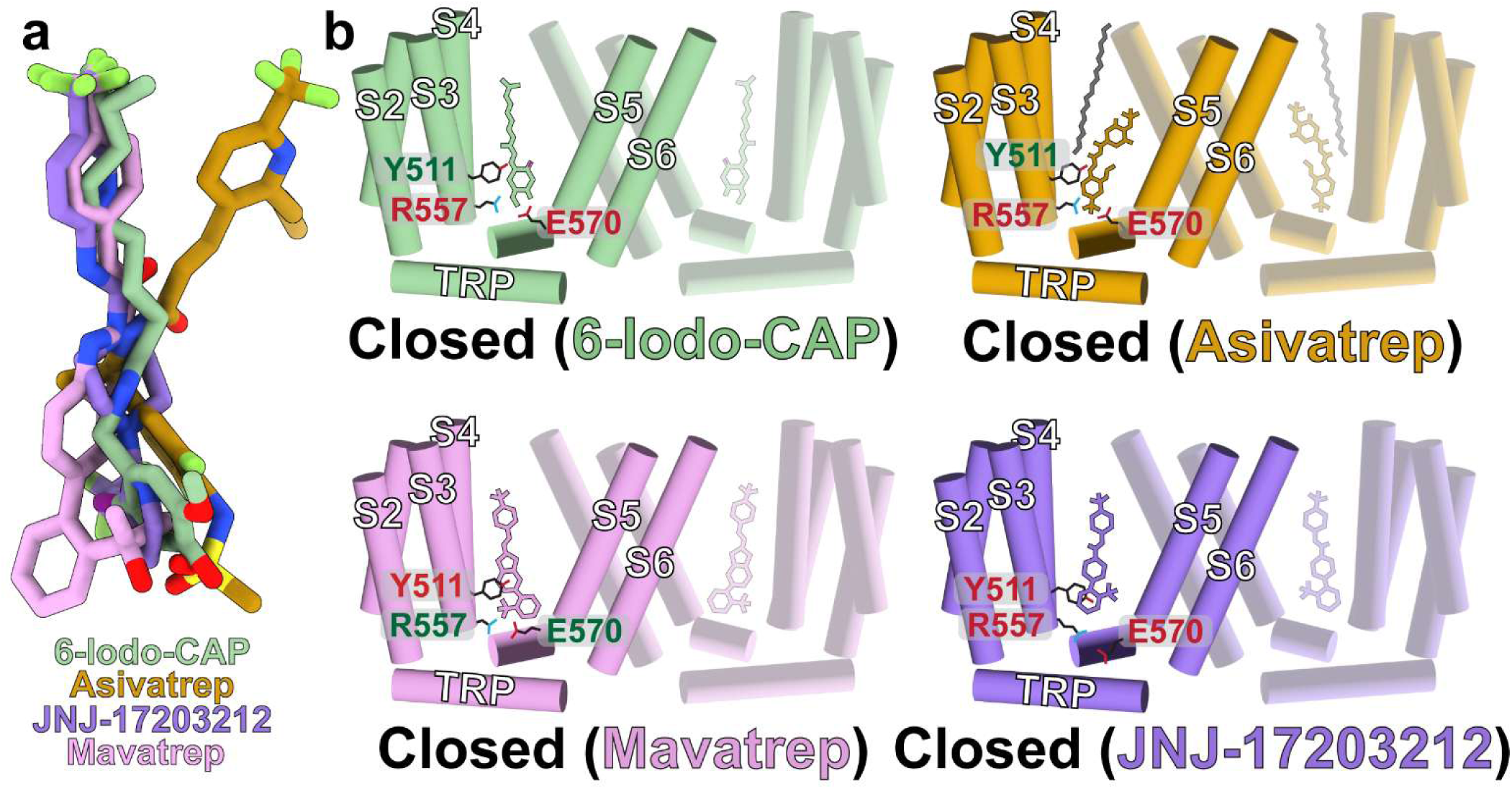
Distinct vanilloid pocket interactions converge on a unified TRPV1 closed state. **a** Overlay of ligands in our structures with structures aligned at the transmembrane domain (residues 416-712). **b** Cartoons for hTRPV1_6-Iodo-CAP_ (green), hTRPV1_Asivatrep_ (orange), hTRPV1_Mavatrep_ (pink), hTRPV1_JNJ-17203212_ (purple) where red labels indicate no interaction and green a strong interaction.

The 6-Iodo-CAP, JNJ-17203212, and Mavatrep tail groups occupy similar poses within the upper vanilloid pocket, whereas the Asivatrep tail is displaced by ∼3.8 Å and the vacated space is occupied by a nominal lipid acyl chain (Fig. 5a-d, 6d, e). This observation demonstrates that the vanilloid pocket can accommodate multiple hydrophobic molecules simultaneously, consistent with previous observations in the hTRPV1_Libvatrep_ where cholesterol occupies the upper pocket region (Fig. 6f)^28^. These findings suggest that ligand affinity and resulting function can be modulated not only by direct protein-ligand interactions, but also by ternary interactions that include endogenous lipid engagement. In contrast to the tail region, antagonist head groups adopt markedly different positions. Only 6-Iodo-CAP and JNJ-17203212 display closely aligned head groups, whereas Mavatrep and Asivatrep head groups are displaced by ∼5.2 Å and ∼4.0 Å, respectively, relative to JNJ-17203212 (Fig. 10a). The vanilloid pocket accommodates these diverse chemotypes through local conformational rearrangements and preserves the closed pore state.

These distinct binding modalities result in divergent molecular interactions that challenge previously proposed rules for TRPV1 antagonism. Prior structural studies suggest that Y511-ligand engagement and R557/E570 salt bridge disruption are required to inhibit TRPV1 through S4/S5 linker and S6 helix movement towards the pore (Fig 1a)^27, 28, 29, 32^. However, hTRPV1_Mavatrep_ retains an intact R557/E570 salt bridge and is in a closed conformation, which demonstrates disruption of this interaction is not required for antagonism and mutagenesis shows Y511 is not an essential hydrogen bond partner for Mavatrep binding (Fig. 7c, 9d, 10b). Similarly, hTRPV1_JNJ-17203212_ lacks direct interaction between JNJ-17203212 and Y511, which adopts a rotated conformation relative to other ligand-bound structures (Fig 7d, 8c, 9e). Furthermore, Mavatrep and Asivatrep engage T550 through a hydrogen bond network that is mediated by a water molecule, a feature not observed in other ligand-bound TRPV1 structures. In previously reported complexes, T550 typically forms either a direct hydrogen bond or hydrophobic contact with the ligand. Across all antagonist-bound structures, however, the S4-S5 linker is consistently displaced towards the pore (Fig. 3h, 5h, 6d, 7e, 10b). This movement represents a shared structural feature that underlies TRPV1 orthosteric vanilloid binding site inhibition across chemically diverse antagonists (Fig. 10b). Together, these findings demonstrate that chemically diverse antagonists exploit vanilloid pocket plasticity yet converge on a conserved gating mechanism that stabilizes the closed TRPV1 state.

Collectively, these structural data show that the human TRPV1 vanilloid-binding pocket exhibits remarkable plasticity, leveraging conformational selection to accommodate chemically diverse antagonists through divergent molecular interaction networks. Previous evidence indicates that the thermosensing and thermogenic liabilities that have plagued generations of promising TRPV1 compounds can be disentangled from efficacious analgesia^14, 24^. Additionally, the compound-specific shifts in potency we observed when functional assays were shifted from 25 °C to 37 °C (Supplementary Fig. 8) provide functional evidence that these chemically diverse antagonists engage distinct allosteric networks. Because these ligands exhibit differential capacities to inhibit baseline temperature activation, our data provide a combined structural and functional rationale for this divergence.. Coupled with future dynamics studies defining how specific binding networks govern channel allostery, it should be possible to exploit and design next-generation therapeutics that mode-selectively modulate ligand-evoked TRPV1 signaling while preserving its role in thermosensing and thermoregulation.

## Methods

### Cheminformatic-guided Ligand Selection and Chemical Space Analysis

Inspired by Backman et al.^71^, a custom Python (3.13.7) pipeline was generated to evaluate and quantify the chemical diversity of select known TRPV1 ligands. Molecular structures of TRPV1-specific ligands were exported from ChemDraw (25.0.2.7) as SMILES strings. These strings were subsequently processed using the RDKit Python library (2025.3.6) to featurize each molecule as a high-dimensional atom-pair fingerprint, represented as a discrete set of topological atom-pair identifiers. These fingerprints were generated using default parameters, capturing topological distances between atom pairs ranging from 1 to 30 bonds. Each distance was stored as a vector encoding the two atom types and their intervening path length. Pairwise Tanimoto similarity coefficients (*s*) were calculated to evaluate structural relationships and subsequently converted into a dissimilarity matrix (*d*) using the following transformation: *d*_i,j_ = 1 − *s*_i,j_. Dimensionality reduction was performed via classical metric multidimensional scaling (MDS) using eigen-decomposition using the SciPy Python library (1.15.3), projecting the dissimilarity matrix into a two-dimensional Euclidean embedding (classical MDS) for the first two principal coordinates. This analysis is fully deterministic and does not rely on random initialization. The resulting chemical latent space representation was plotted with Matplotlib (3.10.0).

The input SMILES data and Python code are available at: https://github.com/vanhornlab/cheminformatics_python_MDS. The resulting MDS plot (Fig. 1a) helped guide the cryo-EM structural studies to rationally select a structurally diverse subset of orthosteric agonists for cryo-EM characterization, ensuring a broad coverage of known inhibitory chemotypes.

Ligands subjected to structural studies were selected based on mechanistic and clinical relevance, commercial availability, and diversity in the MDS latent space against existing human TRPV1 determined structures.

### hTRPV1 Construct Design for Structural and BRET Studies

The human TRPV1 construct used in these studies encoded the common M315I variant (rs222747), which has a global minor allele frequency (MAF) of ∼27%, but frequencies vary substantially across populations, reaching 52-64% in East Asian populations^72, 73^. For structural studies, the corresponding gene was cloned into the pBTSG vector (Addgene #159420)^72, 73, 74^. The vector contains a C-terminal thermostable GFP (TGP) followed by a 8×-His tag and was modified to add a StrepII tag that follows the 8×-His tag (hTRPV1-TGP-8His-StrepII)^74^. This construct was used for structural studies and validated functionally by whole-cell patch-clamp electrophysiology studies as described below (Supplementary Fig. 13a-c).

Inspired by CalfluxVTN and a TRPV1 BRET study, we designed a Ca^2+^-sensing hTRPV1 BRET construct that encodes mNeonGreen, Troponin C domain, and nano luciferase (mNG-TnC-nLuc) in series^75, 76^. Q5 Site-Directed Mutagenesis (New England Biolabs) was used to mutate residue 69 in mNeonGreen from a tyrosine to an alanine, disrupting the chromophore and creating a negative “dead acceptor” control to evaluate if donor emission bleed-through alters the BRET functional outcomes^77^. All hTRPV1 mutants used in this study were also generated by Q5 site-directed mutagenesis. The mNG-TnC-nLuc gene was ordered from IDT in a pUCIDT vector and transferred to a pEG BacMam vector (Addgene #160451) that encodes hTRPV1 to enable baculovirus (BacMam) transduction of pEG-mNG-TnC-nLuc-hTRPV1 into mammalian 293 cells^78^. This construct was functionally validated by whole-cell patch-clamp electrophysiology studies as described below (Supplementary Fig. 13d-f).

### hTRPV1 Expression and Purification

The pBTSG-hTRPV1-TGP-8His-StrepII vector was transformed into DH10Bac cells to generate bacmid DNA that was then transfected into Sf9 cells and incubated for 96 hrs before harvesting the P1 viral stock. Sf9 cells were transduced with P1 virus and the P2 virus was harvested after 72 hrs.

HEK293S GnTI^-^ (ATCC CRL-3022) suspension cells were grown to 3.5-4.0×10^6^ cells/mL in 800 mL Freestyle media supplemented with 2.5% FBS, 100 U/mL penicillin and 100 U/mL streptomycin maintained at 37 °C and 5% CO_2_ and transduced with 6% final concentration of the P2 viral stock. 24 hr post transduction, 10 mM sodium butyrate was added to the culture, and the temperature was lowered to 30 °C. Cells were harvested by centrifugation at 5000 ×g for 15 min after 72 hrs post-transduction.

The resulting cell pellet was resuspended in 20 mL buffer V (20 mM Tris-HCl pH 8.0 and 150 mM NaCl), flash frozen in liquid nitrogen and stored at -80 °C for later purification.

All purification steps are carried out at 4 °C unless otherwise stated. The frozen cell pellet was thawed on ice and supplemented with 1 mM 2-Mercaptoethanol (ßME), 250 µM pheylmethylsulfonyl fluoride (PMSF), 0.8 µM Aprotinin, 4.7 µM Leupeptin, 2 µM Pepstatin A and 2% glyco-diosgenin (GDN). The lysis was incubated for one hr before Dounce homogenization and clarification of cell debris by centrifugation at 5000 ×g for 10 min. The supernatant was then ultracentrifuged at 180,000 ×g for one hr and the resulting supernatant was incubated with 0.5-1 mL Strep-Tactin XT Sepharose resin (Cytiva) preequilibrated with buffer V supplemented with 1 mM ßME and 0.01% GDN for one hr. The resin was then washed with 10 column volumes of buffer V with 1 mM ßME and 0.01% GDN and then eluted with three column volumes of the same wash buffer supplemented with 100 mM biotin. The eluted fractions were collected, concentrated down to 500 µL in a 100-kDa MWCO Amicon centrifugal concentrator and loaded onto a Superose 6 column run at room temperature. Peak fractions that corresponded to hTRPV1-TGP were pooled and concentrated to ∼1.5 mg/mL in a 100-kDa MWCO Amicon centrifugal filter before cryo-EM grid application. Manual whole-cell patch-clamp electrophysiology data show that the hTRPV1 structural construct with a C-terminal thermostable GFP (TGP)-8His-StrepII tag intact is compatible with ligand (CAP) activation and inhibition (capsazepine) at the orthosteric vanilloid binding site, which is the focus of our structural studies (Supplementary Fig. 13a-c).

### Cryo-EM Grid Preparation and Data Collection

Concentrated hTRPV1-TGP were diluted to 0.75 mg/mL (tetramer) with SEC buffer before it was applied to cryo-EM grids. hTRPV1-TGP was incubated with 140 µM capsaicin (Corden Pharma), 142 µM 6-Iodo-CAP (CAY10448, Cayman Chemical), 140 µM Asivatrep (MedChem Express), 125 µM Mavatrep (MedChem Express), 142 µM JNJ-17203212 (MedChem Express) for 30 min before freezing. After incubation, 3 µL sample was applied to a glow discharged UltrAuFoil 1.2/1.3 (300 mesh) holey gold grid (Electron Microscopy Sciences) and blotted for 3 s after a 15 s wait time on a Mark IV Vitrobot (Thermo Fisher Scientific) maintained at 100% humidity and 4 °C. The blotted grids were plunged into liquid ethane and stored in liquid nitrogen until used for imaging.

The datasets for hTRPV1_6-Iodo-CAP_ and hTRPV1_Mavatrep_ were collected at the Pacific Northwest Cryo-EM Center (PNCC) on a Titan Krios operated at 300 keV and equipped with a K3 direct electron detector with a BioContinuum energy filter and slit width set to 10 eV (Gatan). Movies were acquired with SerialEM at a magnification of 105,000 which corresponds to a pixel size of 0.84 Å/pixel and a defocus range of 1.5-2.5 µm (hTRPV1_Mavatrep_) and 0.8-2.5 µm (hTRPV1_6-Iodo-CAP_). The total exposure time was 1.8 s (hTRPV1_Mavatrep_) and 2.2 s (hTRPV1_6-Iodo-CAP_) across 50 frames for a total dose of ∼50 e^-^/Å^2^ at a dose rate of 18.8 e^-^/pixel/s (hTRPV1_Mavatrep_) and 15.3 e^-^/pixel/s (hTRPV1_6-Iodo-CAP_).

The datasets for hTRPV1_Apo_ and hTRPV1_Asivatrep_ were collected at PNCC on a Titan Krios operated at 300 keV and equipped with a Falcon4i detector with a Selectris X energy filter (Thermo Fisher Scientific) and slit width set to 10 eV. Movies were acquired with EPU at a magnification of 165,000 which corresponds to a 0.73 Å/pixel pixel size and a defocus range of 1.5-2.5 µm. The total exposure time was 4.1 s (hTRPV1_Apo_) and 3.7 s (hTRPV1_Asivatrep_) across 50 frames for a total dose of ∼50 e^-^/Å^2^ at a dose rate of 19.5 e^-^/pixel/s (hTRPV1_Apo_) and 26.1 e^-^/pixel/s (hTRPV1_Asivatrep_).

The datasets for hTRPV1_CAP_ and hTRPV1_JNJ-17203212_ were imaged on a Titan Krios at the Eyring Materials Center (EMC) at Arizona State University (ASU) operated at 300 keV and equipped with a Falcon 4i detector with a Selectris X energy filter (Thermo Fisher Scientific) and a 10 eV slit width.

Movies were acquired with EPU at a magnification of 165,000 which corresponds to a 0.7345 Å/pixel pixel size and a 0.6-2.6 and 0.5-1.5 µm defocus range for hTRPV1_CAP_ and hTRPV1_JNJ-17203212_ respectively. The total exposure times were 5.9 s (hTRPV1_CAP_) and 2.1 s (hTRPV1_JNJ-17203212_) across across 50 frames for 60 e^-^/Å^2^ (hTRPV1_CAP_) and 20 e^-^/Å^2^ (hTRPV1_JNJ-17203212_) total dose at a 5.96 e^-^/pixel/s (hTRPV1_CAP_) and 11.5 e^-^/pixel/s (hTRPV1_JNJ-17203212_) dose rates.

### Cryo-EM Data Processing and Refinement

All data processing and refinement was performed in cryoSPARC v4.7 ^79^. Initially 5,694 (hTRPV1_CAP_), 6,125 (hTRPV1_6-Iodo-CAP_), 4,288 (hTRPV1_JNJ-17203212_), 40,605 (hTRPV1_Asivatrep_), 7,284 (hTRPV1_Mavatrep_), and 15,756 (hTRPV1_Apo_) movies were imported into cryoSPARC and motion corrected with patch motion correction. Patch contrast transfer function (CTF) was performed on the motion corrected exposures and exposures were ignored for further processing if the CTF fit was worse than 6 Å, cryoSPARC relative ice thickness parameter was greater than ∼1.5, and total motion distance was greater than ∼150 Å, which resulted in 3,724 (hTRPV1_CAP_), 5,743 (hTRPV1_6-Iodo-CAP_), 4,196 (hTRPV1_JNJ-17203212_), 38,472 (hTRPV1_Asivatrep_), 6,557 (hTRPV1_Mavatrep_), and 13,352 (hTRPV1_Apo_) exposures. 2D templates were generated with the cryo-EM map for hTRPV1 bound to SB-366791 (EMDB-2998) and those templates served as templates for automated template picking with a 175 Å particle diameter.

Particles picked were inspected and 339,996 (hTRPV1_CAP_), 1,660,569 (hTRPV1_6-Iodo-CAP_), 731,442 (hTRPV1_JNJ-17203212_), 4,879,389 (hTRPV1_Asivatrep_), 2,089,583 (hTRPV1_Mavatrep_), and 1,646,090 (hTRPV1_Apo_) particles were extracted with a box size of 416 pixels 4× Fourier cropped (104 pixels). These particles were then run through 2-3 iterative rounds of reference-free 2D classification which resulted in 136,397 (hTRPV1_CAP_), 155,614 (hTRPV1_6-Iodo-CAP_), 240,047 (hTRPV1_JNJ-17203212_), 1,587,657 (hTRPV1_Asivatrep_), 447,590 (hTRPV1_Mavatrep_), and 340,951 (hTRPV1_Apo_) particles. Ab-initio reconstruction was run with 3 or 4 classes followed by 2-3 iterative rounds of 3 or 4 class heterogeneous refinement which resulted in 47,430 (hTRPV1_CAP_), 75,798 (hTRPV1_6-Iodo-CAP_), 106,008 (hTRPV1_JNJ-17203212_), 476,551 (hTRPV1_Asivatrep_), 198,931 (hTRPV1_Mavatrep_), and 153,457 (hTRPV1_Apo_) final particles. Particles were reextracted without binning and with either a 480 pixel (hTRPV1_6-Iodo-CAP_, hTRPV1_JNJ-17203212_, hTRPV1_Asivatrep_, hTRPV1_Apo_), 448 pixel (hTRPV1_CAP_), or 416 pixel box size (hTRPV1_Mavatrep_), and used for non-uniform refinement with C4 symmetry imposed, defocus refinement and global CTF refinement turned on and minimize over per-particle scale turned on. Resolutions were calculated using the gold standard Fourier Shell Correlation (FSC) of 0.143 and resulted in 2.52 Å for hTRPV1_Apo_, 3.06 Å for hTRPV1_CAP_, 2.90 Å for hTRPV1_6-Iodo-CAP_, 2.34 Å for hTRPV1_JNJ-17203212_, 2.10 Å for hTRPV1_Asivatrep_, and 2.37 Å for hTRPV1_Mavatrep_ and. The local resolution was also estimated with the same gold standard FSC criterion. Cyro-EM density was visualized and assessed for quality in UCSF ChimeraX.

### Model Building

The structure for hTRPV1 bound to SB-366791 (PDB 8GFA) was used as the starting model for hTRPV1_Mavatrep_ and apo hTRPV1 (PDB 8GF8) for hTRPV1_Apo-GDN_^29^. For hTRPV1_6-Iodo-CAP_, hTRPV1_JNJ-17203212_, hTRPV1_Asivatrep_, the hTRPV1_Mavatrep_ structure was used as the starting model. For hTRPV1_CAP_, hTRPV1_6-Iodo-CAP_ structure was used as the starting model. Starting models were fit into the cryo-EM density map in UCSF ChimeraX with the fit in map function^80^. Ligand restraints and PDB files were generated with Phenix eLBOW and manually fit into the EM density with UCSF ChimeraX^80, 81^ Phenix real space refinement was run on the map and starting model with one round of simulated annealing and morphing, and five rounds of minimization_global, rigid_body, adp, local_grid_search, secondary restraints and noncrystallographic symmetry restraints^81^. The refined models were then manually adjust in Coot^82^. Data and model visualization for figures was generated in UCSF ChimeraX^80^. The pore radius was calculated for all structures using HOLE^83^. Ligand Q-scores were calculated using the QScore plugin in ChimeraX developed by Tristan Croll that uses previously reported methods for determining Q-scores^84^.

### Bioluminescent Resonance Energy Transfer (BRET)-based TRPV1 Functional Assays

Baculovirus from the pEG-mNG-TnC-nLuc-hTRPV1 WT vector and all mutants were generated using the same method as pBTSG-hTRPV1-TGP-8His-StrepII (see above). For cell culture and transduction, 293T cells (ATCC CRL-3216) were cultured at 37 °C in phenol red-free (PRF) Dulbecco’s Modified Eagle’s Medium (DMEM) supplemented with 10% Fetal Bovine Serum (FBS), 2 mM L-glutamine, and penicillin-streptomycin (100 U/mL) in the presence of 5% CO_2_. All reagents were obtained from Thermo Fisher Scientific. Cells were seeded in 100 mm treated cell culture dishes (VWR) to achieve 20-30% adherent confluence at the time of transduction. For transduction, 9% v/v baculovirus was added dropwise to the seeded dish followed by 48-hour incubation at 37 °C for 48 hours in the presence of 5% CO_2_. Accutase (Corning) was added to detach the transduced cells and resuspended in supplemented PRF DMEM before seeding in a white 96-well, flat bottom, chimney well luminescence plate (Greiner Bio-One) at 10^5^ cells per well. The 96-well plate was then returned to the incubator overnight prior to the assay.

The BRET assays were performed as previously described with the following modifications^76, 85^. Replicate numbers ranged from 6 to 12 and were collected over a minimum of two separate days, and each replicate was a distinct sample from a separate well of cells. Signal was recorded using a Tecan Spark plate reader with the emission filter for nanoLuciferase set to 430-485 nm and 490-545 nm for mNeonGreen. Measurements were performed while cells were in DPBS (Thermo Fisher Scientific) with 5 µM Coelenterazine h (Nanolight). A baseline BRET signal was recorded for each well before ligand addition and approximately twenty minutes later the BRET signal was re-recorded after ligand addition.

For all antagonist assays, an agonist concentration of 250 nM capsaicin was co-applied in the wells with the antagonist ligand. To ensure the isolation of ligand activation under observation from the possibility of concurrent TRPV1 temperature activation, assays were performed at 25 °C. The nanoLuciferase and mNeonGreen signals were averaged over 24 recordings taken during a 20 second per well scan. To ensure that time-averaged BRET values reliably reflected steady-state signal behavior, ΔBRET_obs_ signal was plotted as a function of time (Supplementary Fig. 14a,b). The BRET signal is the ratio of the mNeonGreen over the nanoLuciferase emission intensity. The reported ΔBRET_obs_ signal was determined by subtracting the pre-ligand baseline BRET ratio from the post-ligand addition BRET ratio.

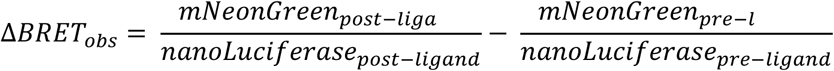

Dose-response curves for both *EC*_50_ and *IC*_50_ (X_50_ in the equation) determination were fitted with a 4-parameter Logistic Equation and plotted with Python 3.12.

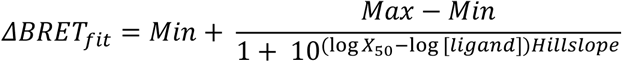

The BRET assays for all ligands were repeated at 37 °C to evaluate how distinct antagonist chemotypes differentially inhibit baseline temperature activation at physiologically relevant temperatures. The assays were performed as previously described and the data processing was also identical to that of the 25 °C assays. Replicate numbers ranged from 6 to 8 and were collected over a minimum of two days. Comparative dose-response plots were made (Supplementary Fig. 8) and a 95% confidence interval was calculated to determine whether shifts in potency were statistically significant.

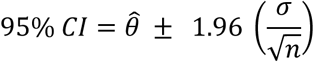

Where *θ̂* is the pEC/IC_50_ fitted parameter, 1.96 is the critical value for a 95% confidence interval, and 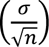 is the standard error of the mean of the pEC/IC_50_.

Baculovirus from the “dead acceptor”, negative control pEG-mNG(Y69A)-TnC-nLuc-hTRPV1 vector was generated using the same method as described previously. BRET assays for capsaicin and Asivatrep were performed at 25 °C using 293T cells transduced with the negative control baculovirus. Four replicates were collected for capsaicin and 3 for Asivatrep. Pre-ligand BRET_mNG_Y69A signal was averaged and subtracted to convert raw basal BRET into net basal BRET and the post-ligand BRET_mNG_Y69A signal was subtracted from the average corresponding BRET_TnC_hTRPV1 post-ligand signal to convert to net post-ligand BRET before calculating ΔBRET_Y69ASub_. ΔBRET_Y69ASub_ was fit with the 4PL equation and comparative dose-response plots were made (Supplementary Fig. 14c-e) As the pEC/IC_50_ and hillslope values were nearly identical, we did not collect any more negative control replicates.

Manual whole-cell patch-clamp electrophysiology confirms the BRET_TnC_hTRPV1 construct functions similarly to hTRPV1 that lacks the Ca^2+^ BRET sensor (mNG-TnC-nLuc) (Supplementary Fig. 13d-f).

### Estimation of Apparent Antagonist Affinities, Mutant Binding Energetics, and Ligand Structure-Activity Thermodynamics

Concentration-response data from cellular BRET assays were fit using a four-parameter logistic model described above. Concentrations were expressed as log_10_ molar concentration, and the fitted midpoint parameter was converted to pIC_50_ or pEC_50_ as *pX_50_ = -log_10_(X_50_)*. The hill coefficient, midpoint, and their standard errors were obtained from the nonlinear regression covariance matrix. To determine the apparent binding affinities (*K_i,app_*) for the TRPV1 antagonists across the wild-type and mutant constructs, we applied the Cheng–Prusoff equation for competitive inhibition. The BRET antagonism assay utilized a fixed capsaicin agonist concentration ([capsaicin]) of 250 nM. The *K_i,app_* was calculated as:

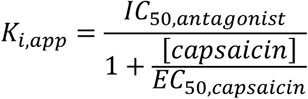

The capsaicin EC_50_ used in the correction was determined independently for each TRPV1 construct. Thus, the WT capsaicin EC_50_ was used for the WT data, whereas corresponding mutant capsaicin EC_50_ was used for each mutant. For agonists (capsaicin and dihydrocapsaicin), the EC_50_ values were treated as apparent dissociation constants (K_d,app_) representing the macroscopic binding affinity. These values should be interpreted as apparent cellular affinities because they are derived from the functional BRET concentration-response data and depend on the assumptions of the competitive inhibition model. Uncertainties in pIC_50_ and pEC_50_ were treated as independent, and first-order Taylor-series propagation was used. Covariance between the antagonist and capsaicin fits was not included because the fits were performed independently:

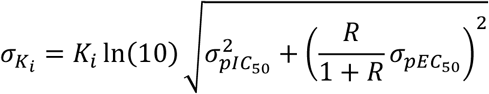

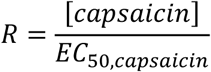

To evaluate the energetic impact of individual mutations or ligand structural modifications, changes in the apparent binding free energy (ΔΔG) was calculated relative to the wild-type construct. Values were calculated at 298.15 K (25 °C) where RT ≈ 0.592 kcal/mol.

For evaluating ligand modification (e.g., saturation of the aliphatic tail or halogenation), apparent standard free energies of binding (ΔG°) were calculated for each ligand using their respective affinities (*K* = *K*_i,app_ for antagonists and *K* = *EC*_50_ for agonists):

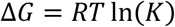

The thermodynamic shift induced by the chemical modification was then calculated as the difference between the modified ligand and the parent scaffold:

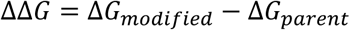

For evaluating receptor mutations, using the ratio of the mutant and wild-type apparent binding affinities, the change in the Gibbs free energy can be estimated by:

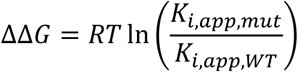

Positive values indicate weaker apparent antagonist binding to the mutant relative to wild-type, whereas negative values indicate stronger apparent binding. For each antagonist, ΔΔG was calculated relative to wild-type construct for the same antagonist. Wild-type values were therefore assigned ΔΔG = 0 by definition. Similarly, for ligand modifications, positive values denote an energetic penalty associated with the functional group modification.

The uncertainty for the thermodynamic shift (σ_ΔΔG_) was propagated from the absolute errors of both the mutant and the wild-type *K_i,app_* values using standard methods for independent variables:

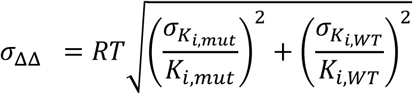

The compared affinity estimates were treated as independent. Reported uncertainties for *K_i,app_* and ΔΔG are propagated fitting uncertainties derived from the covariance-based standard errors of the antagonist and capsaicin nonlinear regressions. They do not include additional uncertainty from assay-to-assay biological variation, model misspecification, or uncertainty in the fixed capsaicin concentration.

Fits were inspected individually for convergence, parameter plausibility, and adequate coverage of the response transition. Constructs for which the concentration-response curve did not support a reliable midpoint estimate (EC_50_/IC_50_) were excluded from the Cheng–Prusoff and ΔΔG analyses. The outcomes of the mutational analyses are shown in Table 1.

### Whole-cell Patch-clamp Electrophysiology Validation of Cryo-EM and BRET Constructs

293 cells (ATCC CRL-1573) were cultured at 37 °C in DMEM (Thermo Fisher) supplemented with Premium Plus Fetal Bovine Serum (Gibco), Penicillin-Streptomycin (Cytiva), and L-Glutamine (Thermo Fisher). For pBTSG-hTRPV1-TGP-8His-StrepII functional verification, transient transfection of 293 cells with 500 ng of the expression plasmid and FuGENE 6 (Promega). TGP epifluorescence was used to guide cells expressing the hTRPV1-TGP-8His-StrepII fusion protein. For the BRET_TnC_hTRPV1 functional verification, 293T cells were transduced by addition of P2 baculovirus (9% v/v).

The extracellular solution (ECS) contained 132 mM NaCl, 4.8 mM KCl, 1.2 mM MgCl_2_ hexahydrate, 2 mM CaCl_2_ dihydrate, 10 mM HEPES, 5 mM Glucose, pH adjusted to 7.4 using 1M NaOH. The ECS osmolality was adjusted to 311 mOsm using sucrose as measured by vapor pressure osmometry (Wescor Vapro 5600). The intracellular solution (ICS) contained 135 mM potassium gluconate, 5 mM KCl, 1 mM MgCl_2_, 5 mM EGTA, 10 mM HEPES, 2 mM sodium ATP, pH of the ICS was adjusted to 7.2 using 1M KOH and the osmolality was adjusted to 300 mOsm with sucrose.

Pipettes were pulled from borosilicate glass capillary tubes (World Precision Instruments) using a P-2000 laser puller (Sutter Instruments) Tips were heat polished with an MF-830 micro forge (Narashige) and pipette resistance was 2-5 MΩ. A 2% agar and intracellular solution bridge held the ground electrode. All experiments were performed at 25 °C.

Whole-cell voltage-clamp measurements were collected following established protocols with a Molecular Devices Axopatch 200B amplifier and Axon Digidata 1550B using Clampex 10.7 software^16^. Seven replicates were collected over a period of two days for pBTSG-hTRPV1-TGP-8His-StrepII functional verification. Gravity fed perfusion was used to apply 500 nM capsaicin to the cells resting on glass coverslips in the extracellular solution to evaluate agonism, 500 nM capsaicin and 500 nM capsazepine were subsequently co applied to evaluate antagonism. Five replicates were collected over a period of two days for BRET_TnC_hTRPV1 functional verification. Gravity fed perfusion was used to apply 250 nM capsaicin to evaluate agonism, 250 nM capsaicin and 500 nM capsazepine were subsequently co-applied to evaluate antagonism. All electrophysiology experiments used a voltage-clamp pulse program where initial potential is held at of -80 mV followed by applied voltage steps between -120 mV and +160mV. The current response for each step at the 300 ms recording time point was plotted using SigmaPlot 12.0 to create a current-voltage (IV) plot to show capsaicin agonism and capsazepine antagonism at each membrane potential step.

## Data Availability

All data needed to evaluate the conclusions of the paper are present in the paper or the supplementary information. The cryo-EM density maps were deposited to the Electron Microscopy Data Bank (EMDB) under the accession codes EMD-75616 (hTRPV1_Apo_), EMD-78595 (hTRPV1_CAP_) EMD-75617 (hTRPV1_6-Iodo-CAP_), EMD-75619 (hTRPV1_Mavatrep_), EMD-75618 (hTRPV1_Asivatrep_), and EMD-78331 (hTRPV1_JNJ-17203212_). The atomic coordinates were deposited too the Protein Data Bank (PDB) under the accession codes 11CJ (hTRPV1_Apo_), 37XE (hTRPV1_CAP_) 11CK (hTRPV1_6-Iodo-CAP_), 11CN (hTRPV1_Mavatrep_), 11CL (hTRPV1_Asivatrep_), and 37NU (hTRPV1_JNJ-17203212_). All other data are available from the corresponding author upon request.

## Acknowledgments

We acknowledge the support from the National Institute of Neurological Disorders and Stroke R01NS119505 (W.D.V.H.); National Institutes of General Medical Sciences R35GM141933 (W.D.V.H.); The American Heart Association 23POST1029540 (K.E.L.); Arizona State University School of Molecular Sciences Presidential Postdoctoral fellowship (K.E.L.). A portion of this research was performed in the John Cowley Center for High Resolution Transmission Electron Microscopy at Arizona State University on a Krios G2 funded by the National Science Foundation Major Instrumentation grant: MRI 1531991. Images were collected on a Selectris X imaging filter and Falcon 4i combination funded by the National Institutes of Health’s Shared Instrumentation Grant, SIG 1S10OD036204-01, with assistance from Dr. Dewight Williams. A portion of this research was supported by NIH grant R24GM154185 and performed at the Pacific Northwest Center for Cryo-EM (PNCC) with assistance from Drs. Marcello De Farias and Marzia Miletto. We thank Dr. Samrat Amin in the Magnetic Resonance Research Center (MRRC) for setting up and maintaining computational resources. We thank Sadie Heeringa for cloning the modified pBTSG vector and Drs. Helen Bank and Dustin Luu for cloning the BRET TRPV1 construct into a BacMam vector. The content is solely the responsibility of the authors and does not necessarily represent the official views of any funding agencies.

## Contributions

W.D.V.H. designed the study. K.E.L. carried out protein expression, purification, cryo-EM data collection, processing, model building and analysis. A.S.P. carried out BRET assays and electrophysiology experiments. M.D. carried out MDS chemoinformatic analysis. K.E.L. and W.D.V.H. wrote the manuscript. All authors edited the manuscript.

