## Supplemental Files for "Chemically diverse antagonists inhibit human TRPV1 through ligand-specific interaction networks"

**Contents:**

**Supplementary Figures S1-S9 and Table S1**

*Supplementary Figure 1: hTRPV1_Apo_ cryo-EM data processing, collection, and validation*

*Supplementary Figure 2: hTRPV1_CAP_ cryo-EM data processing, collection, and validation*

*Supplementary Figure 3: Structural comparisons of hTRPV1_Apo_, hTRPV1_CAP_, and rTRPV1_Open_*

*Supplementary Figure 4: hTRPV1_6-Iodo-CAP_ cryo-EM data collection, processing, and validation*

*Supplementary Figure 5: hTRPV1_JNJ-17203212_, hTRPV1_Asivatrep_, hTRPV1_Mavatrep_ cryo-EM data collection, processing and validation*

*Supplementary Figure 6: Vanilloid pocket plasticity for hTRPV1_SAF312_, hTRPV1_SB-366791_, and hTRPV1_CAP_*

*Supplementary Figure 7: Pore analysis for hTRPV1 antagonist-bound cryo-EM structures*

*Supplementary Figure 8: Functional BRET assay comparison at 25 °C and 37 °C*

*Supplementary Figure 9: Cryo-EM structural analysis of hTRPV1_Asivatrep_*

*Supplementary Figure 10: Cryo-EM structural analysis of hTRPV1_Mavatrep_*

*Supplementary Figure 11: Cryo-EM structural analysis of hTRPV1_JNJ-17203212_*

*Supplementary Figure 12: Functional effects of hTRPV1 mutations on capsaicin and antagonist responses*

*Supplementary Figure 13: Functional characterization of TRPV1 constructs*

*Supplementary Figure 14: BRET assay validation and background subtraction*

*Supplementary Table 1: Molecular structures, pharmacological behavior, and MDS coordinates of TRPV1 ligands*

*Supplementary Table 2: Cryo-EM data collection, refinement and validation statistics*

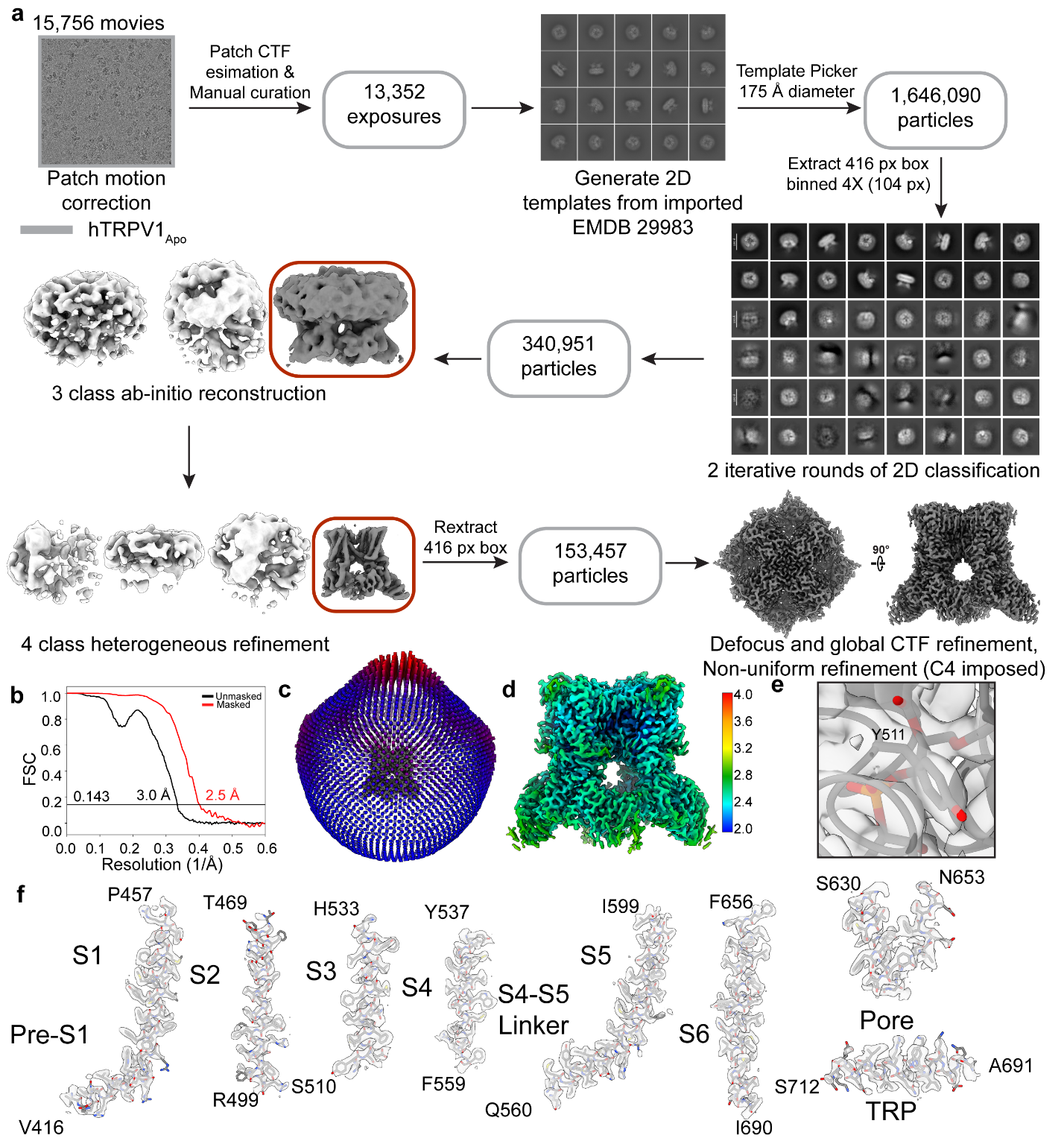

**Supplementary Figure 1: hTRPV1_Apo_ cryo-EM data processing collection and validation. a** Data collection and processing workflow for hTRPV1_Apo_. **b** Fourier shell correlation for the final map refinement with masked (red) and unmasked (black) curves displayed and the resolution using the gold standard cutoff (0.143) marked with a horizontal black line. **c** Euler angle distribution plot. **d** Local resolution estimates. **e** Cryo-EM density for Y511 where the residue and PI lipid are shown as sticks. **f** Map and model for each transmembrane helix with every residue represented as sticks.

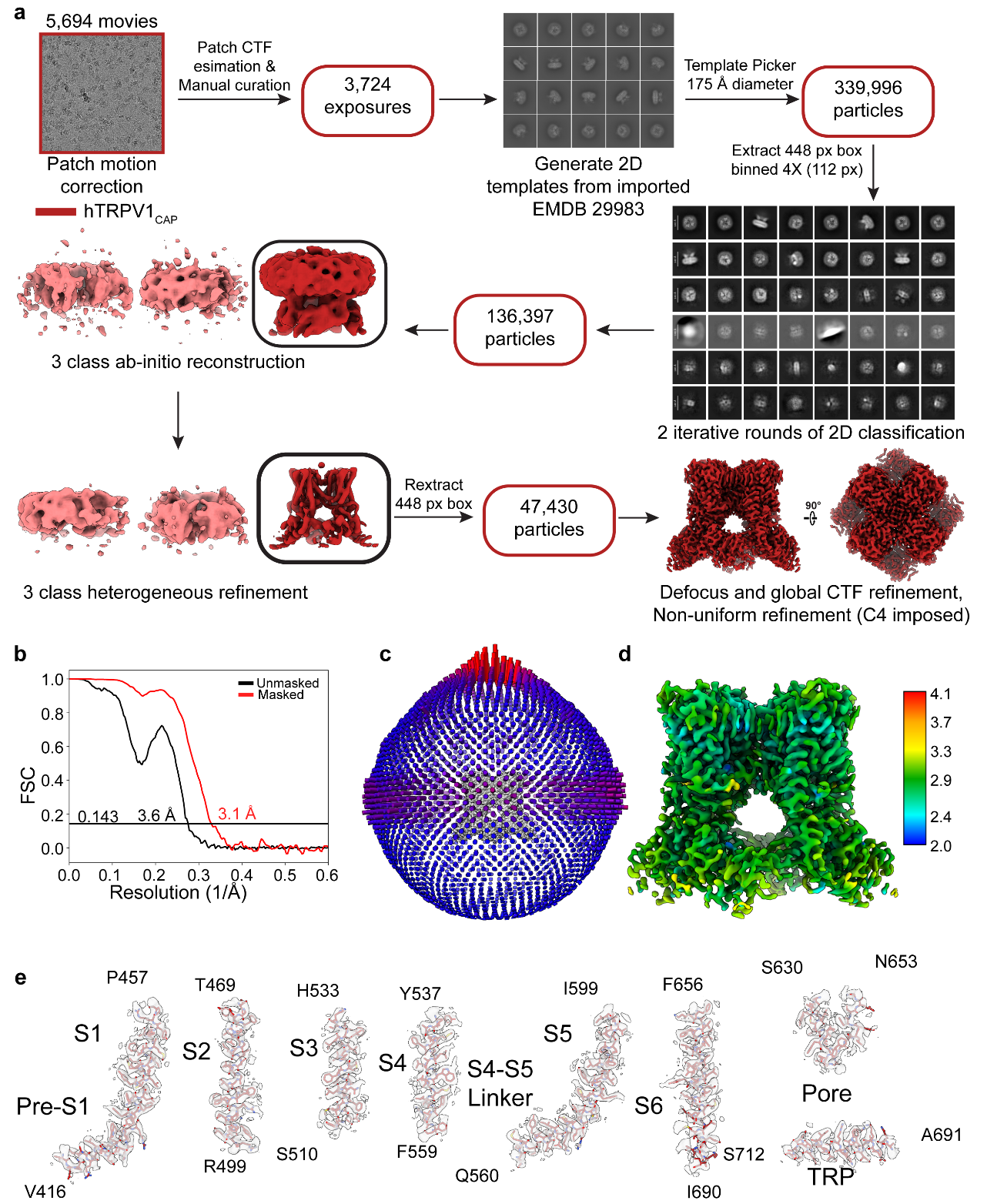

**Supplementary Figure 2: hTRPV1_CAP_ cryo-EM data processing collection and validation. a** Data collection and processing workflow for hTRPV1_Apo_. **b** Fourier shell correlation for the final map refinement with masked (red) and unmasked (black) curves displayed and the resolution using the gold standard cutoff (0.143) marked with a horizontal black line. **c** Euler angle distribution plot. **d** Local resolution estimates. **e** Map and model for each transmembrane helix with every residue represented as sticks.

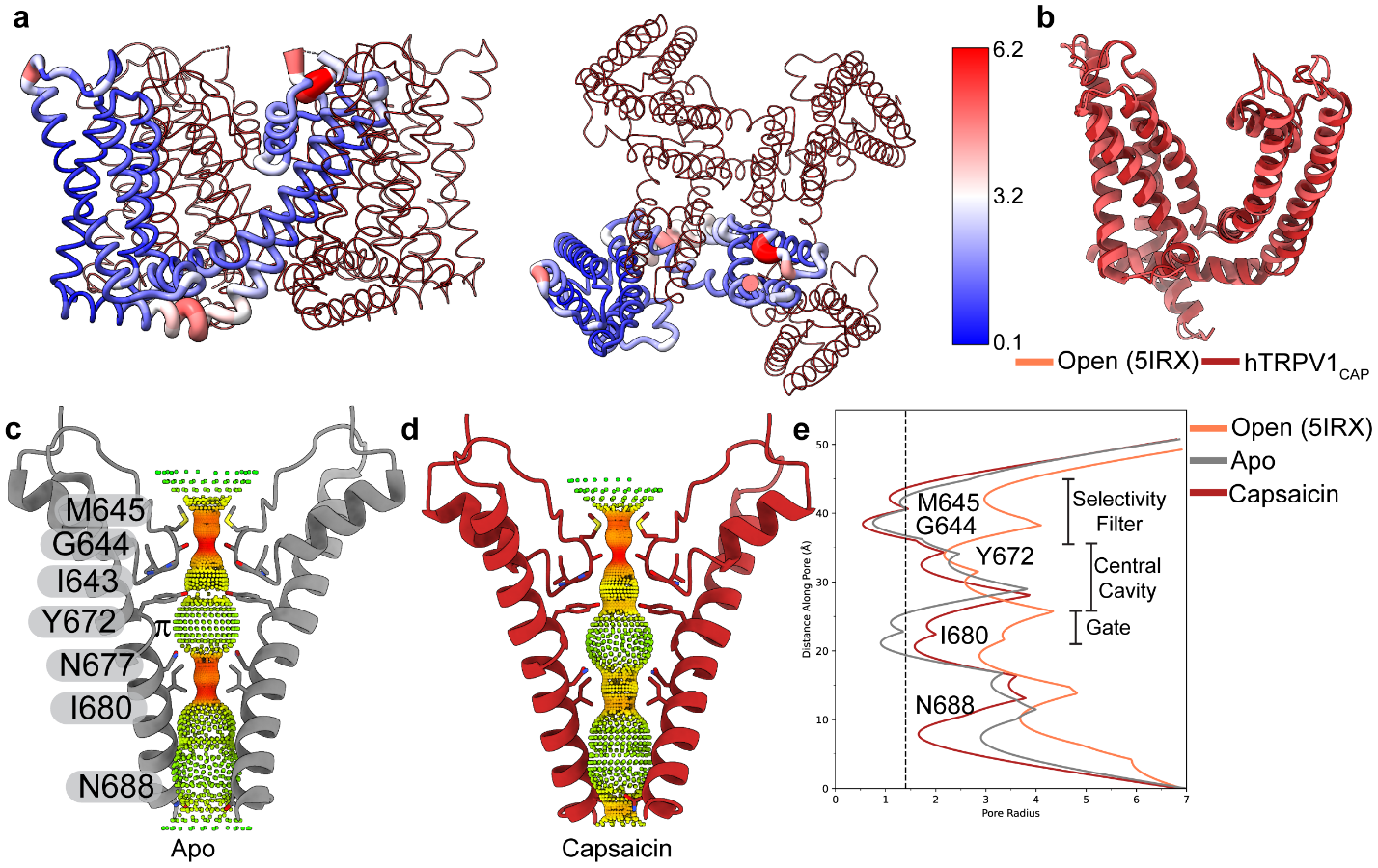

**Supplementary Figure 3: Structural comparisons of hTRPV1_Apo_, hTRPV1_CAP_, and rTRPV1_Open_. a** hTRPV1_CAP_ model is displayed showing C_α_ RMSD between hTRPV1_CAP_ and rTRPV1_Open_ (PDB 5IRX), determined by alignment of residues 416 to 712. Large changes are indicated by wider, red tubes and small to no change by smaller, blue tubes (scale bar to the right). **b** Same overlay as in **a**. Both hTRPV1_CAP_ (red) and rTRPV1_Open_ (coral) models shown to show the extent of the changes. HOLE analysis for hTRPV1_Apo_ (**c**) and hTRPV1_CAP_ (**d**) where radius size colored red (widest), yellow (intermediate) and green (narrowest) and pore lining residues are displayed and labeled. **e** Plot for HOLE analysis for hTRPV1_CAP_ (red), hTRPV1_Apo_ (grey), and the rTRPV1_Open_ (coral; PDB 5IRX). Pore lining residues are labeled, and the radius of a water molecule is represented by a dashed line (1.4 Å).

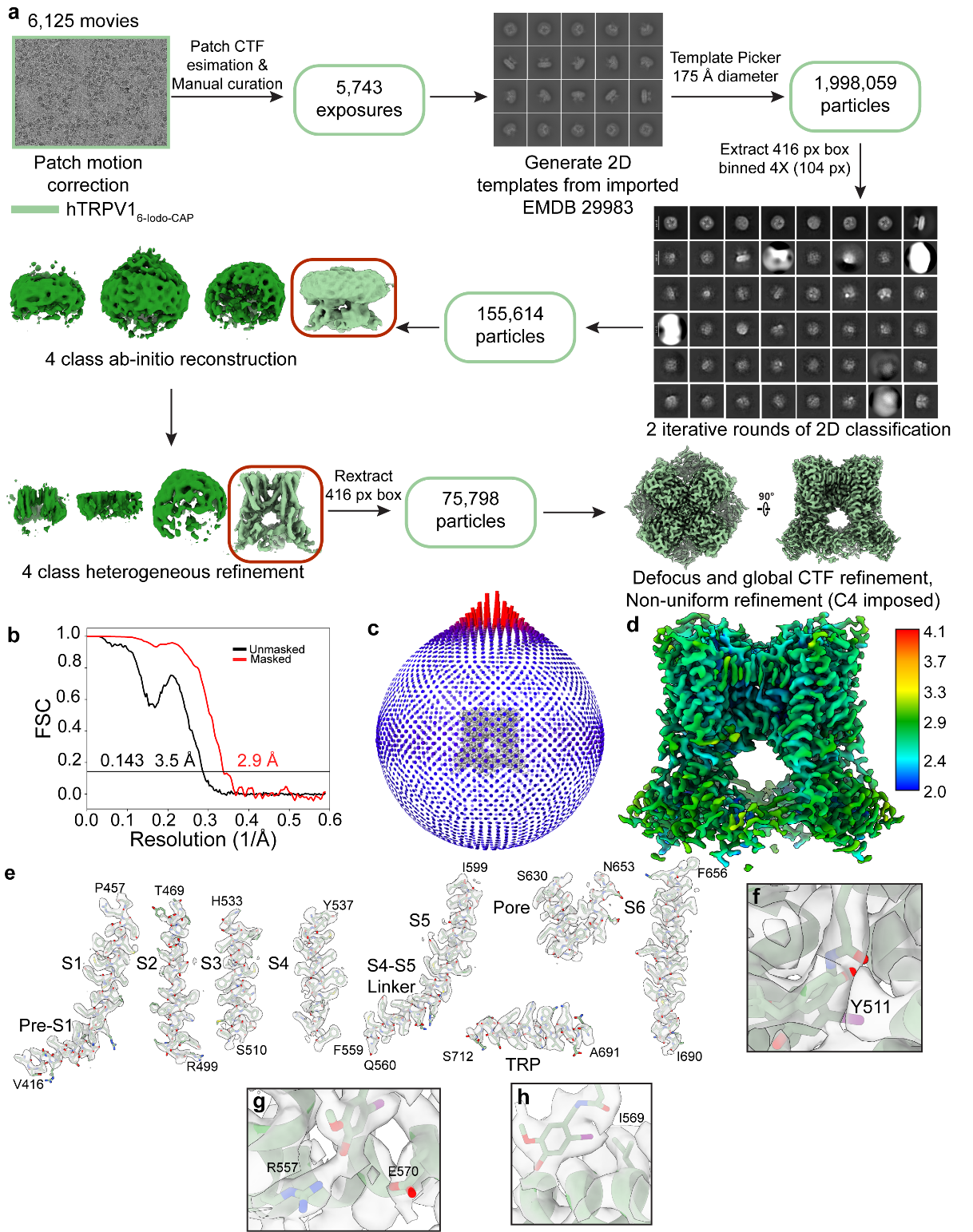

**Supplementary Figure 4: hTRPV1_6-Iodo-CAP_ cryo-EM data collection, processing, and validation. a** Data collection and processing workflow for hTRPV1_6-Iodo-CAP_. **b** Fourier shell correlation for the final map refinement with masked (red) and unmasked (black) curves displayed and the resolution using the gold standard cutoff (0.143) marked with a horizontal black line. **c** Euler angle distribution plot. **d** Local resolution estimate. **e** Map and model for each transmembrane helix with every residue represented as sticks. Density for residues Y511 (**f**), R557 and E570 (**g**), and I569 (**i**) with residues labeled and represented as sticks.

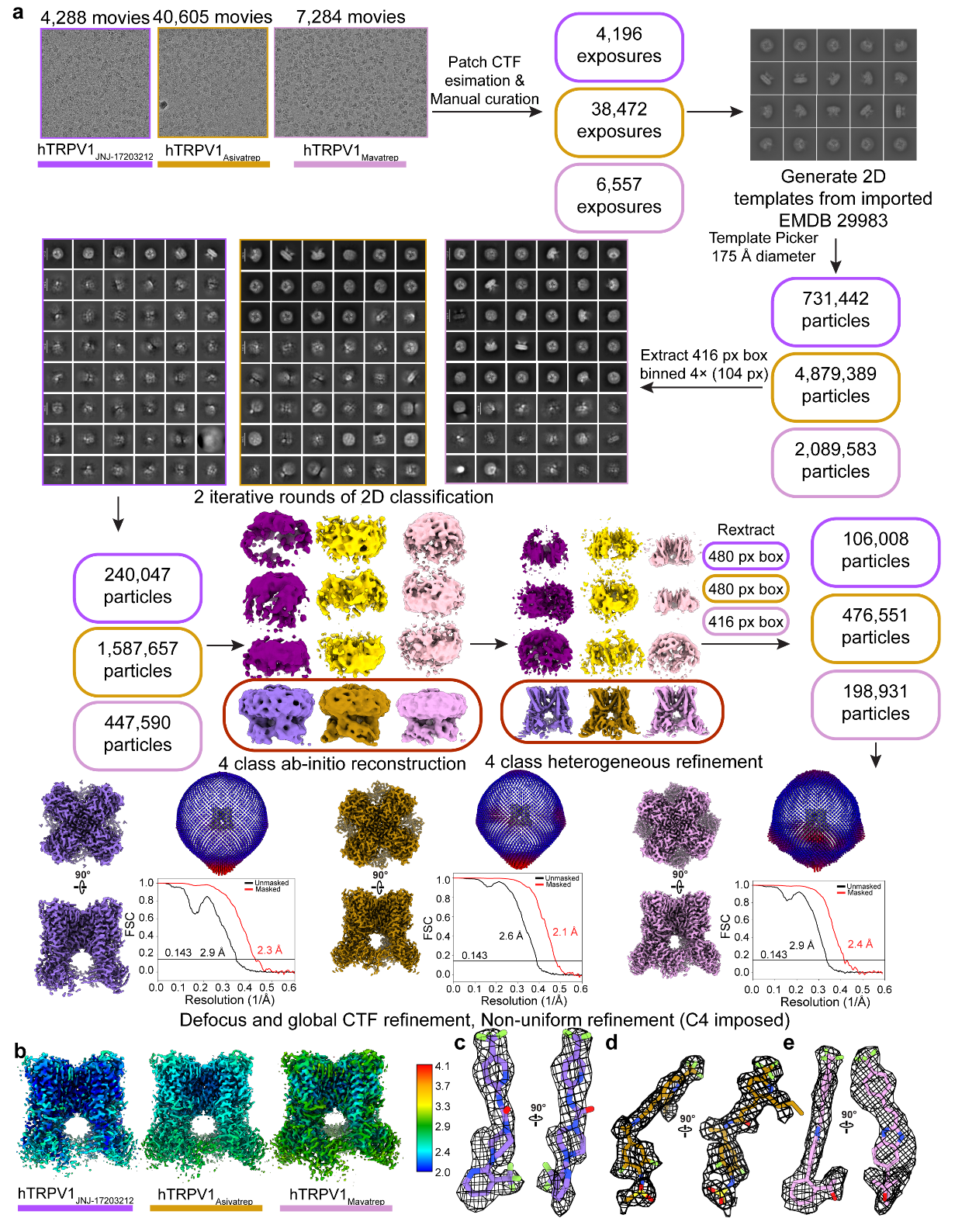

**Supplementary Figure 5: hTRPV1_JNJ-17203212_, hTRPV1_Asivatrep_, hTRPV1_Mavatrep_ cryo-EM data collection, processing and validation. a** Data collection and processing workflow for hTRPV1_JNJ-17203212_ (purple), hTRPV1_Asivatrep_ (orange, and hTRPV1_Mavatrep_ (pink). Cryo-EM map top and bottom views displayed in corresponding colors, Euler angle distribution plots and FSC plots for masked (red) and unmasked (black) displayed and gold standard resolution cutoff (0.143) shown. **b** Local resolution estimates for hTRPV1_JNJ-17203212_, hTRPV1_Asivatrep_, and hTRPV1_Mavatrep_ with labels for each structure underneath. Ligand model and density for JNJ-17203212 (**c**), Asivatrep (**d**), and Mavatrep (**e**).

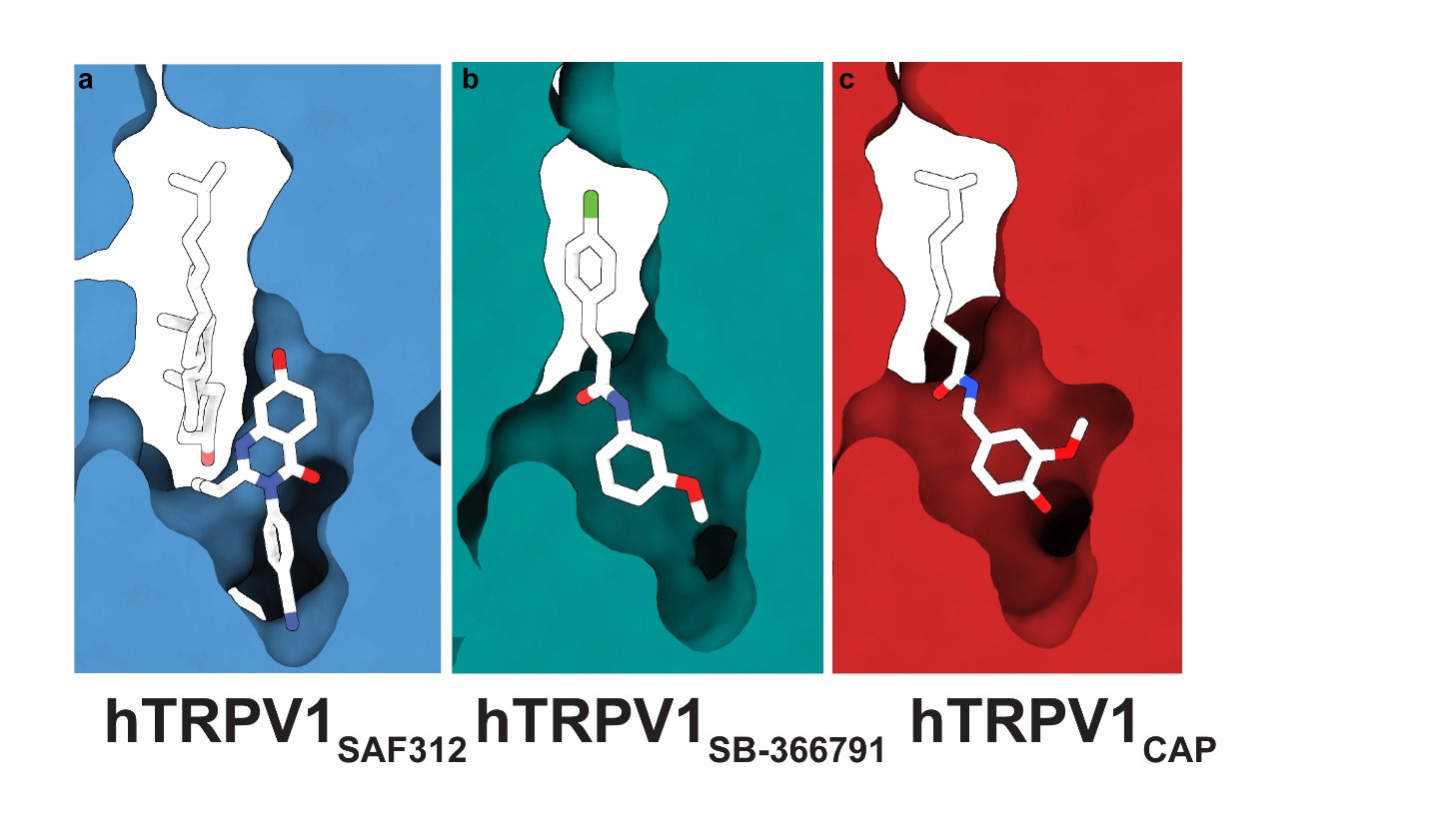

**Supplementary Figure 6: Vanilloid pocket plasticity for hTRPV1_SAF312_, hTRPV1_SB-366791_, and hTRPV1_CAP_.** Experimental models for hTRPV1_SAF312_ (PDB 8JQR, **a**), hTRPV1_SB-366791_ (PDB 8GFA, **b**), and hTRPV1_CAP_ (**c**) represented as a surface viewed from inside-out the vanilloid pocket. The ligand models are displayed as white sticks.

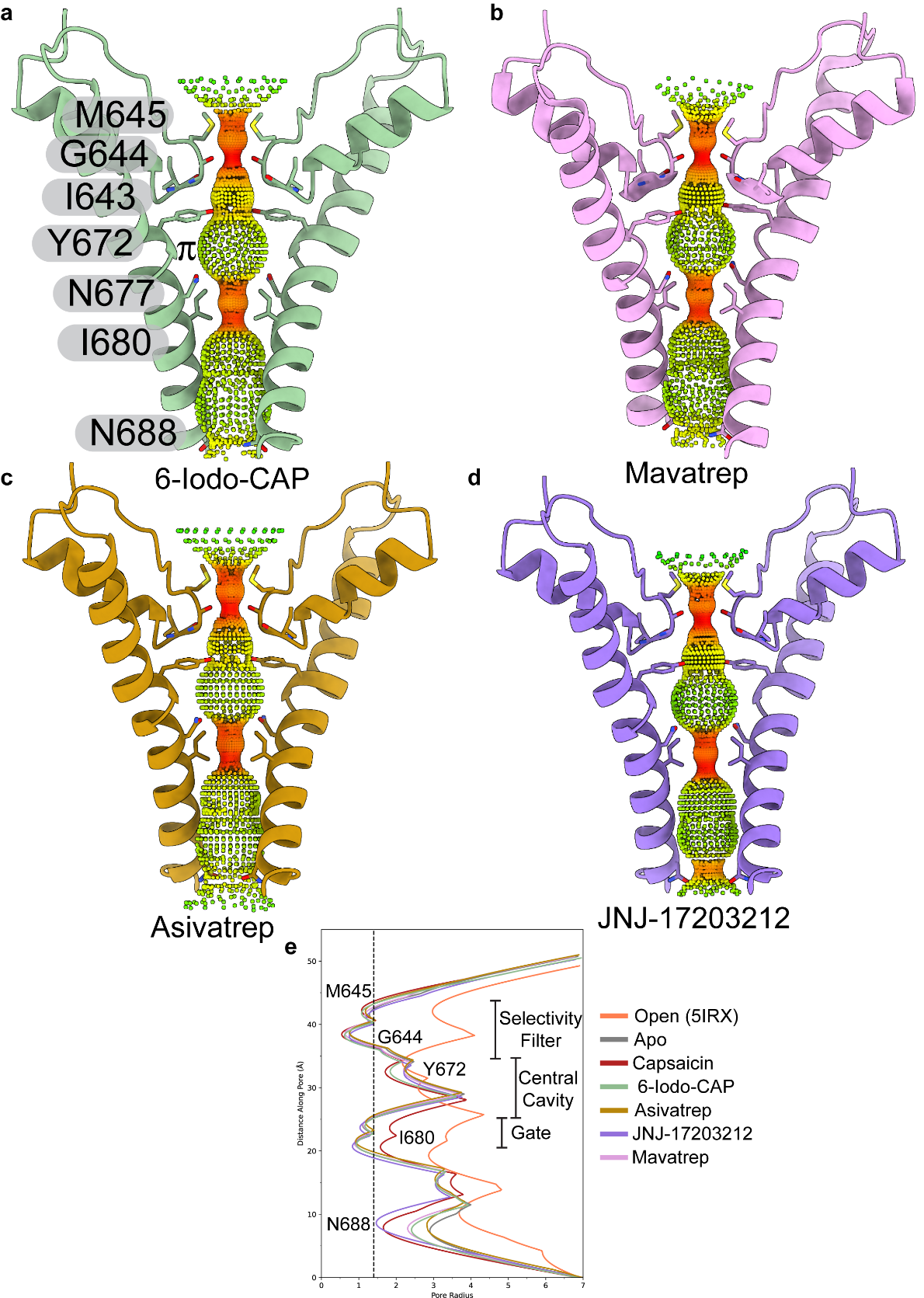

**Supplementary Figure 7: Pore analysis for cryo-EM structures.** HOLE analysis where radius size colored red (widest), yellow (intermediate) and green (narrowest) and pore lining residues are displayed and labeled for hTRPV1_6-Iodo-CAP_ (**a**), hTRPV1_Mavatrep_ (**b**), hTRPV1_Asivatrep_ (**c**), and hTRPV1_JNJ-17203212_ (**d**). **e** Plot for HOLE analysis for our structures and rTRPV1_Open_ (red; PDB 5IRX). Pore lining residues are labeled, and the radius of a water molecule is represented by a dashed line (1.4 Å).

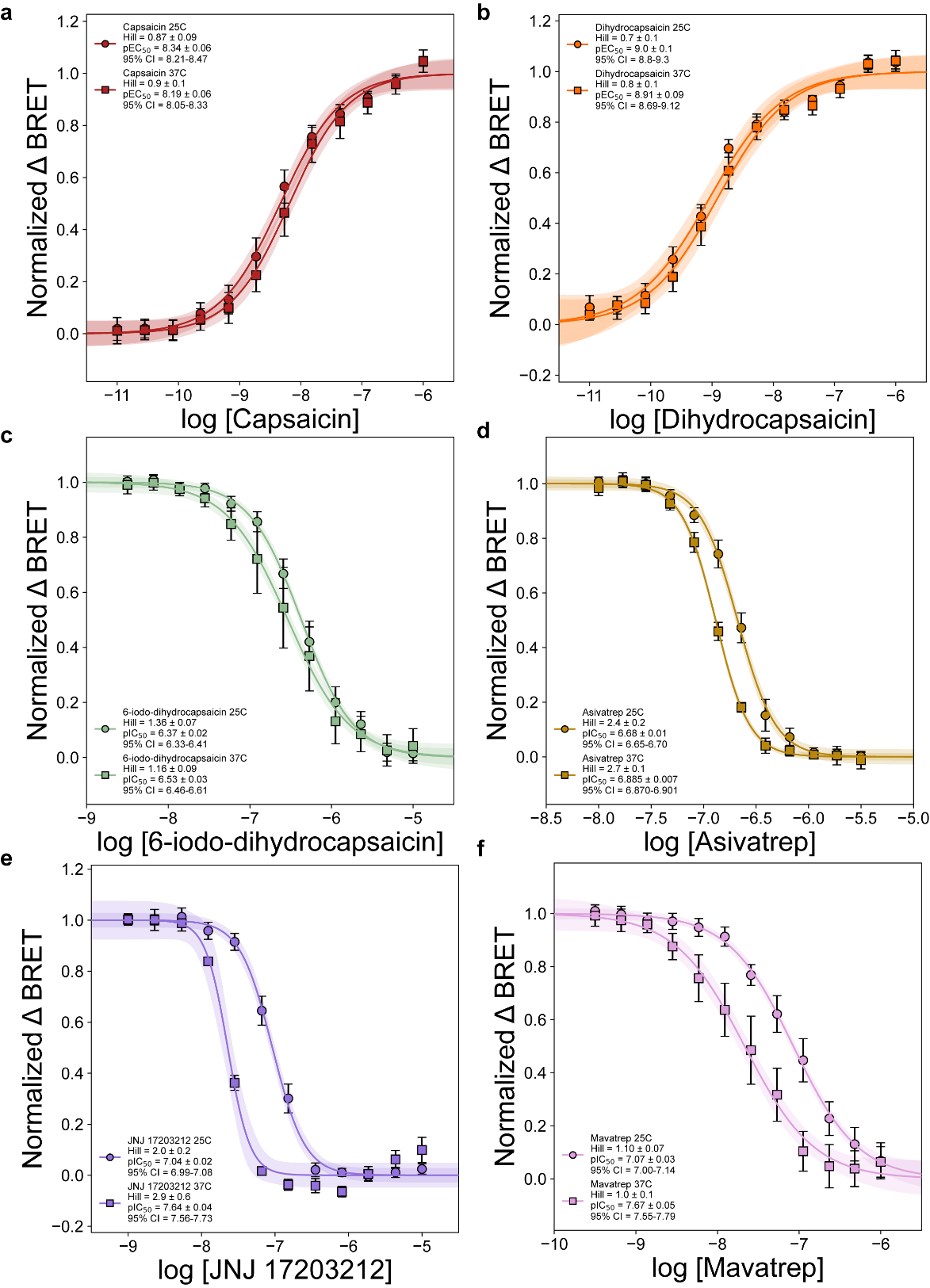

**Supplementary Figure 8: BRET experiments compared at 25 °C and 37 °C.** ΔBRET dose response curves for capsaicin (**a**), dihydrocapsaicin (**b**), 6-Iodo-CAP (**c**), Asivatrep (**d**), JNJ-17203212 (**e**), and Mavatrep (**f**) where data were recorded at 25 °C (cirlces) and 37 °C (squares) and antagonist measurements were performed in the presence of 250 nM capsaicin. Hill error is shown as the SEM and pEC/IC_50_ error is shown by both the SEM as well as the 95% confidence interval (CI) for a fitted parameter.

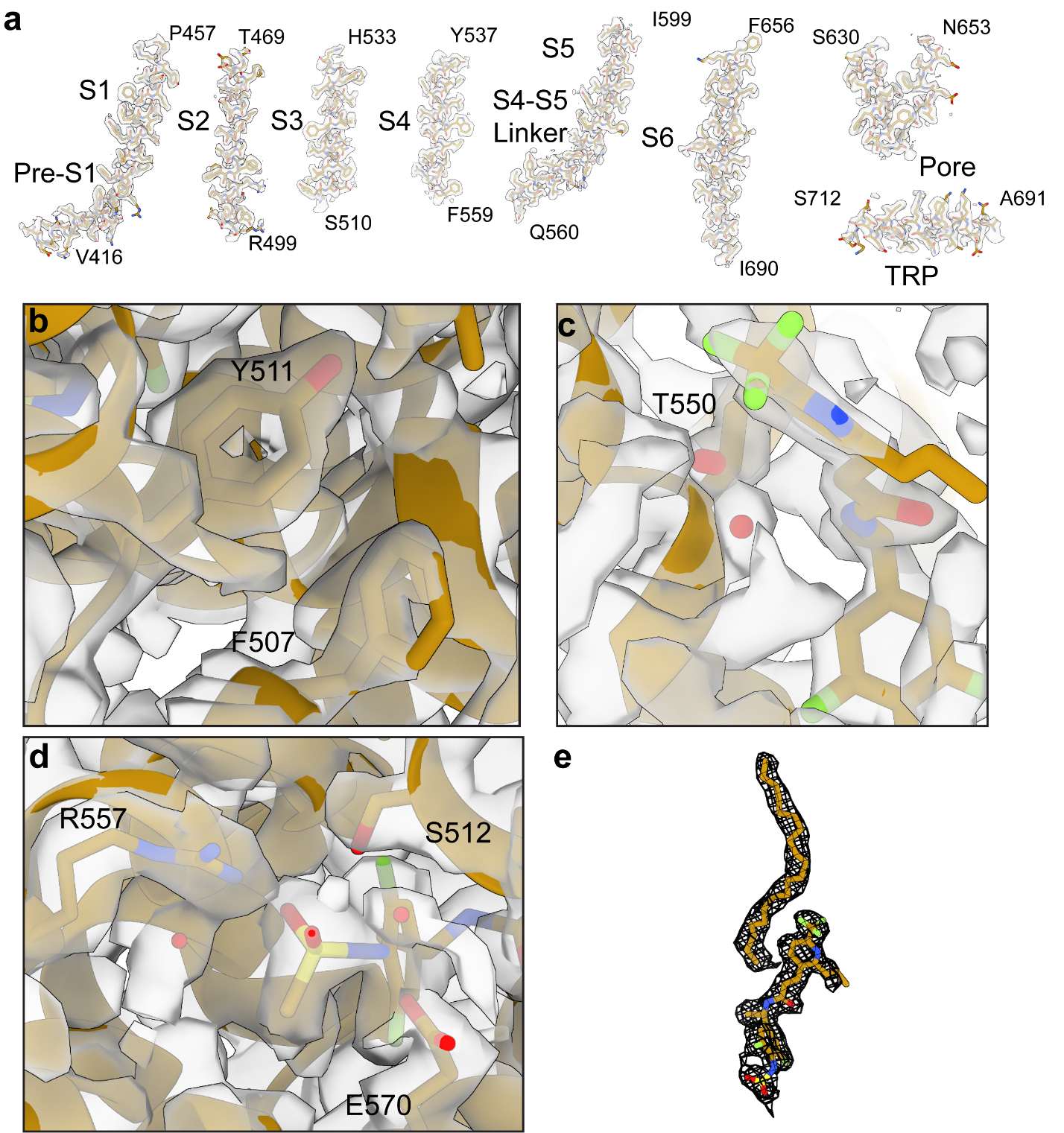

**Supplementary Figure 9: Cryo-EM structural analysis of hTRPV1_Asivatrep_. a** Map and model for each transmembrane helix with every residue represented as sticks. Cryo-EM densities for Y511 and F507 (**b**), T550 and bound water molecule (**c**), and R557, S512, E570 and bound water molecules (**d**). **e** Cryo-EM density (black mesh) and model (orange) for Asivatrep ligand (bottom) and eicosane molecule (top).

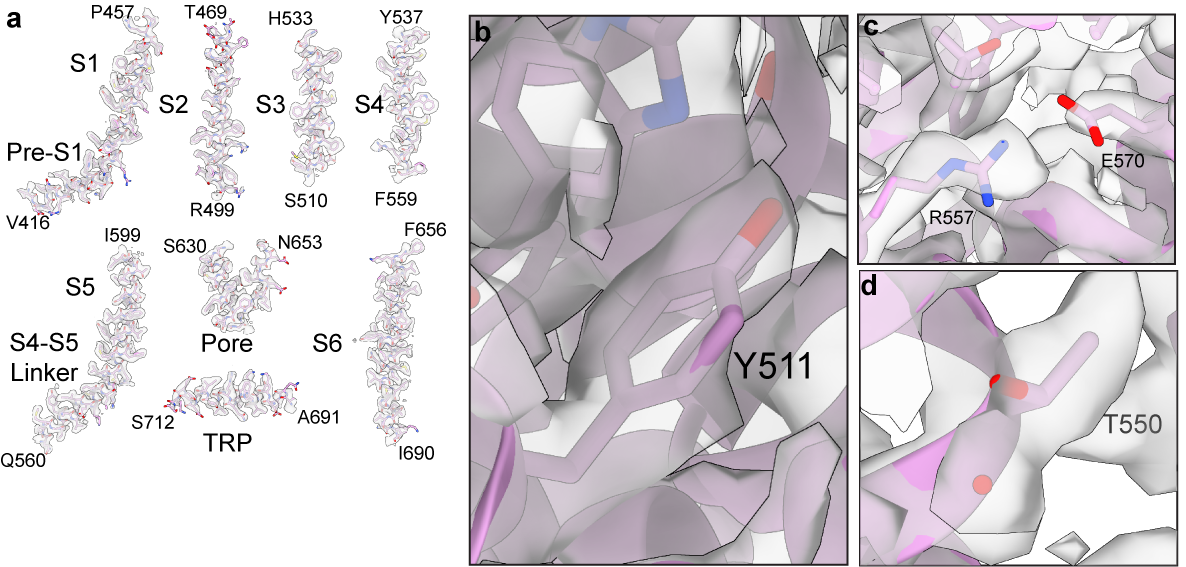

**Supplementary Figure 10: Cryo-EM structural analysis of hTRPV1_Mavatrep_. a** Map and model for each transmembrane helix with every residue represented as sticks. Cryo-EM density and model shown for Y511 (**b**), R557 and E570 (**c**), and T550 and a water molecule (**d**).

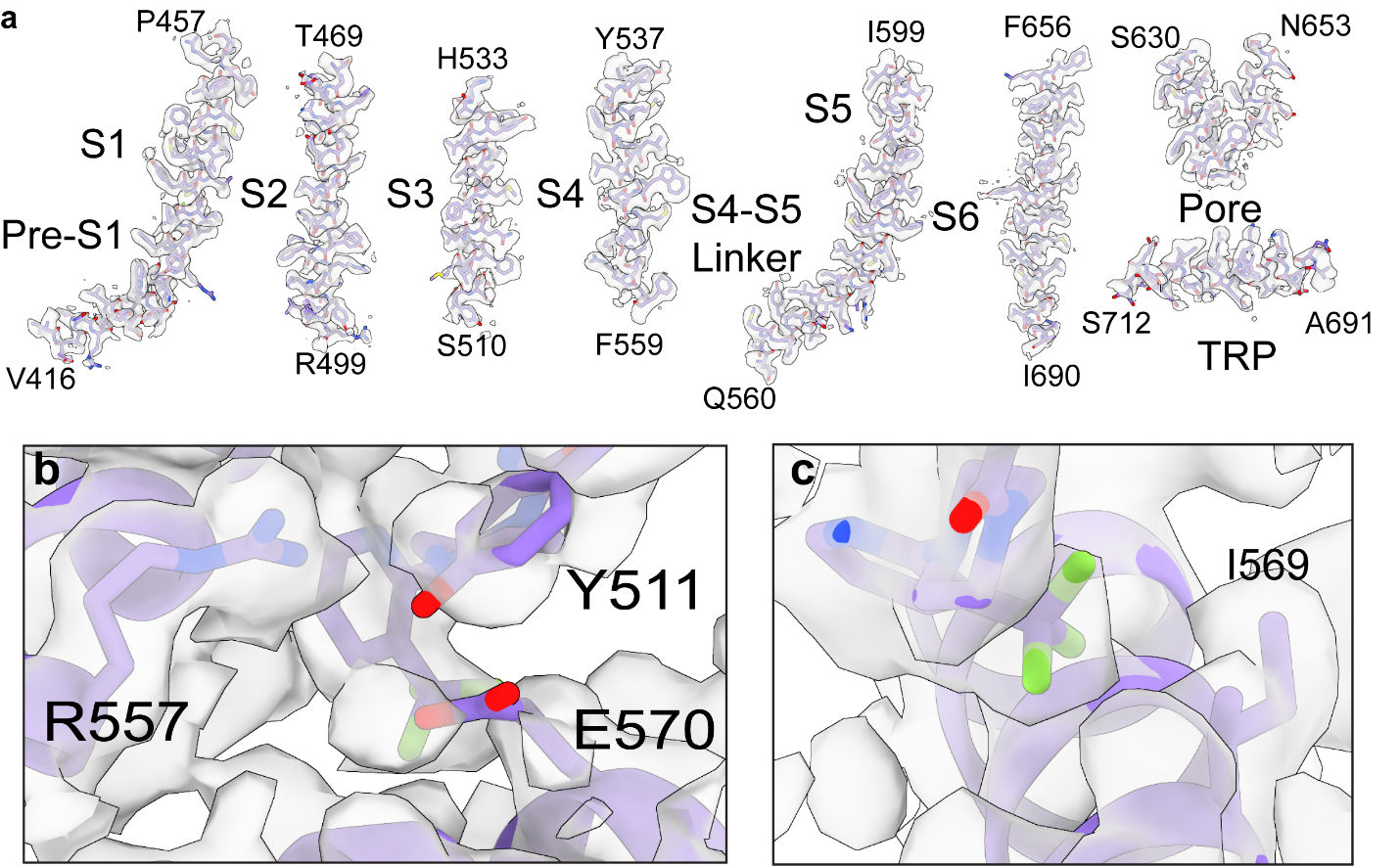

**Supplementary Figure 11: Cryo-EM structural analysis of hTRPV1_JNJ-17203212_. a** Map and model for each transmembrane helix with every residue represented as sticks. Cryo-EM density and model shown for R557, E570, and Y511 (**b**) and I569 (**c**).

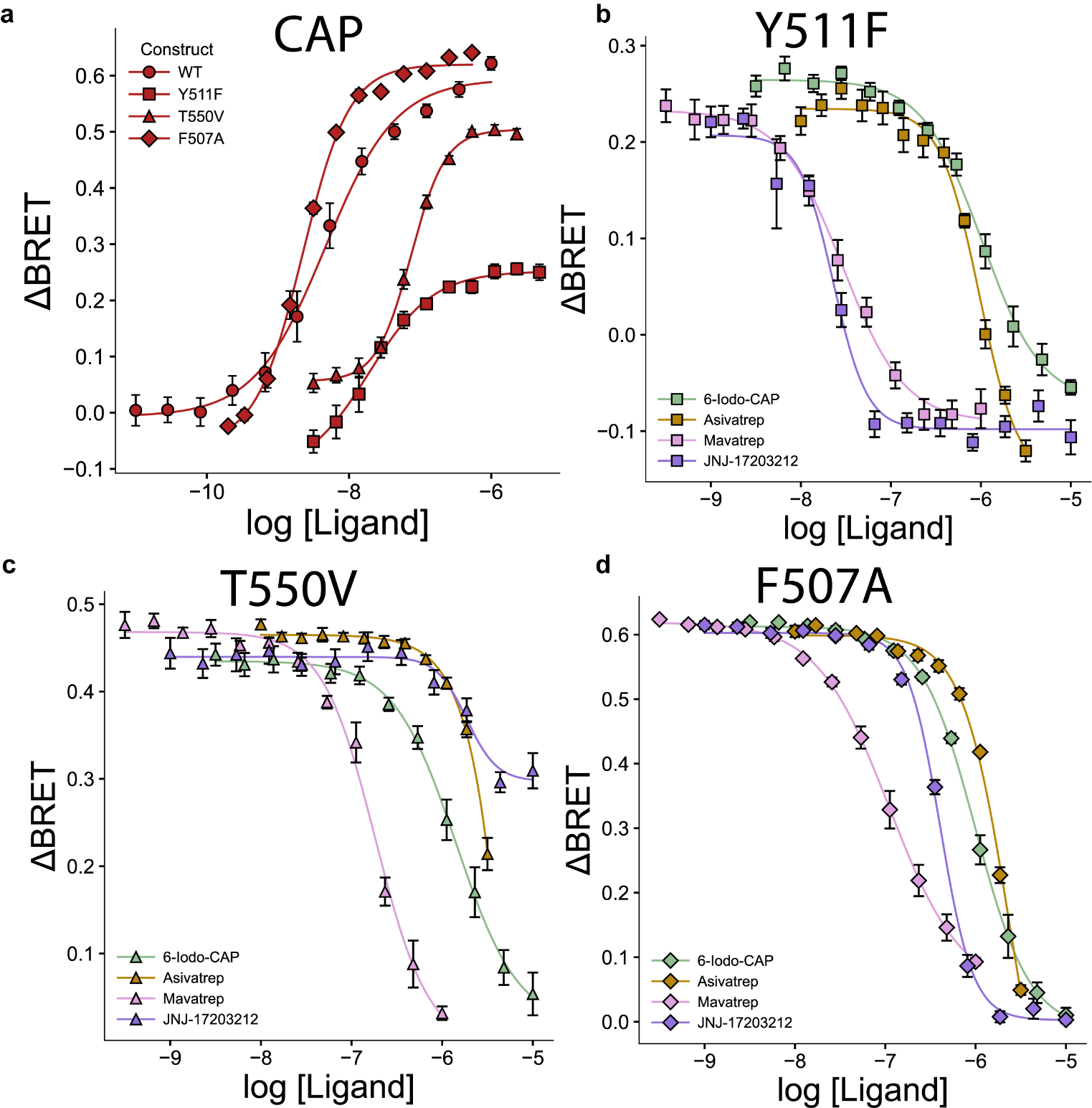

**Supplementary Figure 12: Functional effects of hTRPV1 mutations on capsaicin and antagonist responses. a** Dose response BRET curves for capsaicin with wild-type (WT; circles), Y511F (squares), T550V (triangles), and F507A (diamonds) curves overlayed. Dose response BRET curves for 6-Iodo-CAP (green), Asivatrep (orange), Mavatrep (pink), JNJ-17203212 (purple) overlayed for the mutants Y511F (**b**), T550V (**c**), and F507A (**d**) in the presence of 250 nM capsaicin. The error bars are SEM and the pIC_50_/pEC_50_ and hill coefficient values are derived from fitting errors.

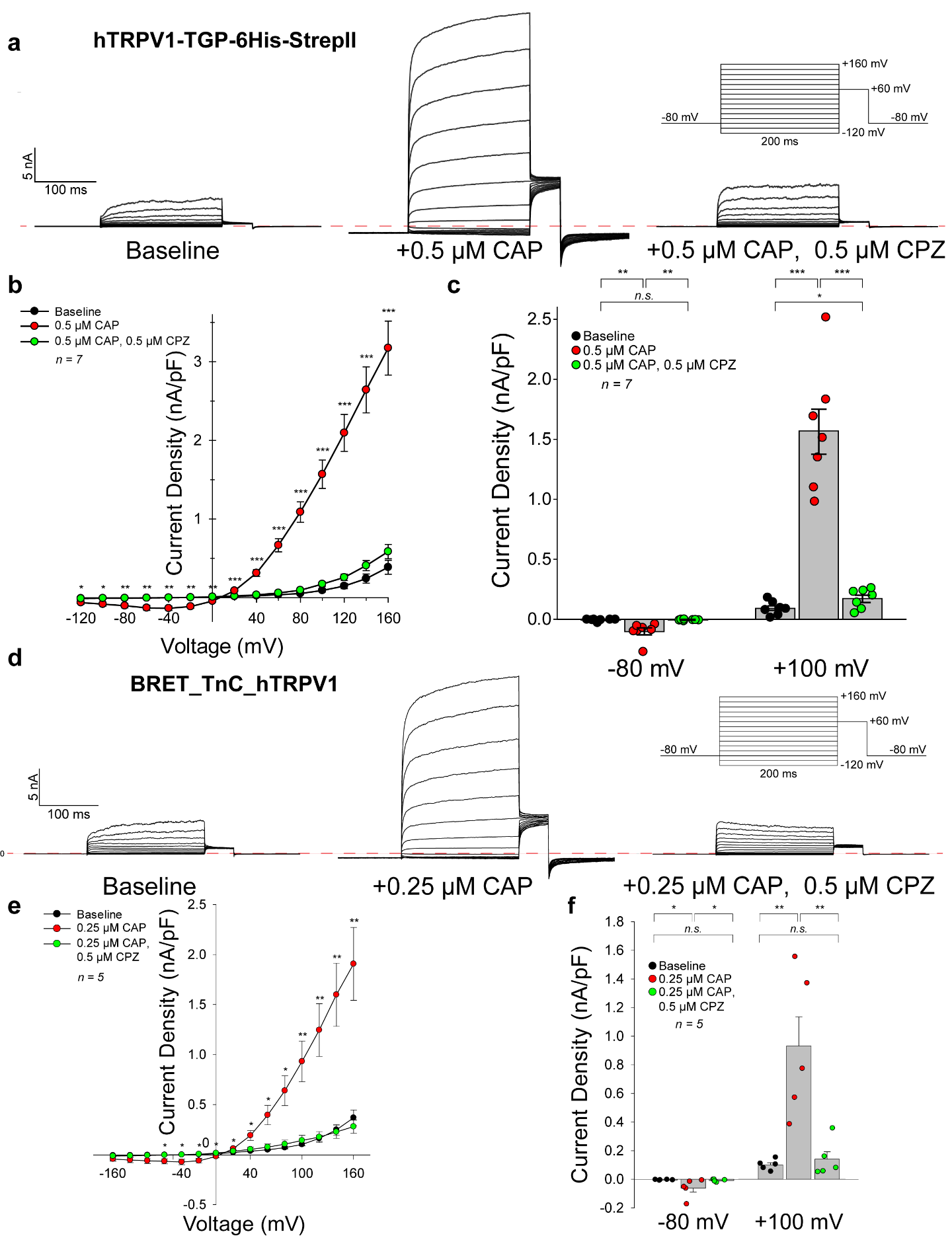

**Supplementary Figure 13: Functional characterization of TRPV1 constructs. a** Averaged (n = 7) whole-cell manual patch-clamp recordings for structural construct hTRPV1-TGP-6His-StrepII using a voltage-clamp step protocol from -120 mV to +160 mV in 20 mV increments. Sweeps show baseline current, current response during perfusion of 500 nM capsaicin (CAP), and current response during subsequent co-application of 500 nM CAP and 500 nM capsazepine (CPZ). **b** Current-voltage (IV) plot generated from the sweeps shows averaged current density at the 300 ms time point for each of the voltage steps under baseline conditions (black), CAP activation (red), and CAP + CPZ co-application (green). Error is the standard error of the mean. Statistical significance was determined using a one-sided paired student’s t-test to compare the CAP and CAP + CPZ current response. P ≤ 0.05 (*), P ≤ 0.01 (**), P ≤ 0.001 (***). **c** Bar plots show the averaged current density at the -80 mV and +100 mV voltage steps with individual replicates jittered to show the spread of cell current responses. Error is the standard error of the mean. **d** Averaged (n = 5) whole-cell manual patch-clamp recordings for BRET construct BRET_TnC_hTRPV1 using a voltage-clamp strep protocol from -120 mV to +160 mV in 20 mV increments. Sweeps show baseline current, current response during perfusion of 250 nM CAP and 500 nM CPZ. **e** Current-voltage (IV) plot generated from the sweeps shows averaged current density at the 300 ms time point for each of the voltage steps under baseline conditions (black), CAP activation (red), and CAP + CPZ co-application (green). Error is the standard error of the mean. Statistical significance was determined in the same way as (**b**). **f** Bar plots show the averaged current density at the -60 mV and +100 mV voltage steps with individual replicates jittered to show the spread of cell current response. Error is the standard error of the mean.

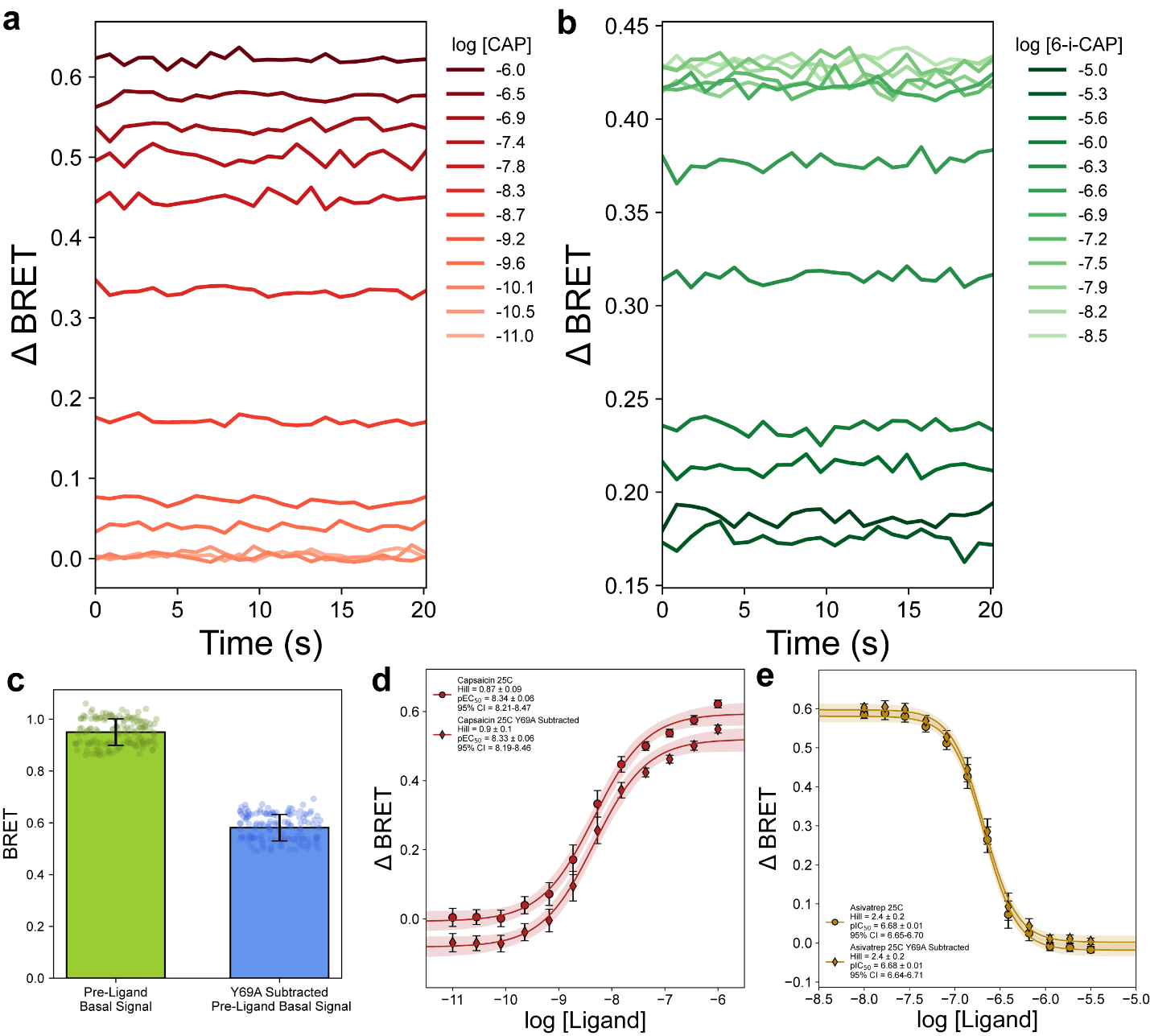

**Supplementary Figure 14: BRET assay validation and background subtraction.** ΔBRET signal over a 20 s time course with increasing concentrations of CAP (**a**) and 6-Iodo-CAP (**b**). Traces represent the average of 12 replicates per concentration for both CAP and 6-Iodo-CAP. **c** Basal BRET signal before ligand addition for the wild-type hTRPV1 BRET construct (left, green, n = 144) and the Y69A mNeonGreen dead hTRPV1 BRET construct (right, blue, n = 48). Individual replicates are jittered and error bars represent the standard deviation. ΔBRET dose response curves for capsaicin (**d**) and Asivatrep (**e**). Circles represent data calculated without Y69A subtraction and diamonds represent data with Y69A subtraction. Error bars represent the standard error of the mean (SEM) and error on fit is shown with a 95% confidence interval (CI). Asivatrep data points were measured in the presence of 250 nM capsaicin.

**Supplementary Table 1: Molecular structures, pharmacological behavior, and MDS coordinates of TRPV1 ligands**

| **Compound** | **SMILES** | **Behavior** | **MDS1** | **MDS2** |
| --- | --- | --- | --- | --- |
| Capsaicin | OC1=C(OC)C=C(CNC(CCCC/C=C/C(C)C)=O)C=C1 | Agonist | -0.447 | 0.079 |
| Dihydrocapsaicin | O=C(NCC1=CC=C(C(OC)=C1)O)CCCCCCC(C)C | Agonist | -0.510 | 0.109 |
| A-1165442 | O=C(NC1=CC=CC2=C1C=C(C)N=C2)N[C@H]3C(C=CC(Cl)=C4)=C4OC(CF)(CF)C3 | Antagonist | 0.050 | -0.105 |
| A425619 | O=C(NCC1=CC=C(C(F)(F)F)C=C1)CC2=CC=CC3=CN=CC=C32 | Antagonist | 0.187 | -0.002 |
| A-784168 | O=C(NC1=CC=C(S(=O)(C(F)(F)F)=O)C=C1)C(CC2)=CCN2C3=C(C(F)(F)F)C=CC=N3 | Antagonist | 0.244 | 0.252 |
| ABT-102 | CC(C)(C1=CC(CC[C@@H]2NC(NC3=C4C=NNC4=CC=C3)=O)=C2C=C1)C | Antagonist | 0.038 | -0.036 |
| AMG-517 | O=C(C)CC1=NC2=C(OC3=CC(C4=CC=C(C(F)(F)F)C=C4)=NC=N3)C=CC=C2S1 | Antagonist | 0.146 | -0.091 |
| AMG-7905 | FC(F)(F)C(C=C1)=CC(NCC2CCCCC2)=C1C3=NC=NC(NC4=CC=C(N=CS5)C5=C4)=C3 | Antagonist | 0.181 | 0.110 |
| AMG 9810 | O=C(/C=C/C1=CC=C(C(C)(C)C)C=C1)NC2=CC=C(OCCO3)C3=C2 | Antagonist | -0.074 | -0.151 |
| CAY10448 | O=C(NCC1=C(I)C=C(C(OC)=C1)O)CCCCCCC(C)C | Antagonist | -0.509 | 0.108 |
| Capsazepine | OC1=C(O)C=C2C(CN(C(NCCC3=CC=C(Cl)C=C3)=S)CCC2)=C1 | Antagonist | -0.101 | 0.093 |
| JNJ-17203212 | O=C(NC1=NC=C(C(F)(F)F)C=C1)N(CC2)CCN2C3=NC=CC=C3C(F)(F)F | Antagonist | 0.280 | 0.302 |
| JNJ-39439335 (Mavatrep) | FC(F)(F)C(C=C1)=CC=C1/C=C/C2=NC3=CC=C(C4=CC=CC=C4C(C)(O)C)C=C3N2 | Antagonist | 0.192 | -0.144 |
| NEO-6860 | O=C(N[C@H](C1=CC=C(C(C)(C)C#N)C=C1)C)CN2C3=C(C#N)C=CC=C3N=C2 | Antagonist | 0.037 | -0.190 |
| PAC-14028 (Asivatrep) | CCCC1=NC(C(F)(F)F)=CC=C1CCC(N[C@H](C)C2=CC(F)=C(NS(C)(=O)=O)C(F)=C2)=O | Antagonist | 0.013 | 0.099 |
| PHE377 (V-377, PharmEste) | FC(F)(F)C1=CC=C(CNC(NC(C=C2)=CC=C2S(NCC3=CC(CO)=C(O)C=C3I)(=O)=O)=S)C=C1 | Antagonist | 0.034 | 0.096 |
| SB 366791 | O=C(NC1=CC(OC)=CC=C1)/C=C/C2=CC=C(Cl)C=C2 | Antagonist | -0.009 | -0.360 |
| SB 452533 | O=C(NC1=C(Br)C=CC=C1)NCCN(C2=CC=CC(C)=C2)CC | Antagonist | -0.054 | 0.093 |
| SB 705498 | O=C(NC1=CC=CC=C1Br)N[C@@H]2CCN(C3=NC=C(C(F)(F)F)C=C3)C2 | Antagonist | 0.185 | 0.209 |
| V-116517 | O=C(N1CCC(C2=NC=C([C@H](O)CO)C=C2Cl)=CC1)NC3=NC=C(C(F)(F)F)C=C3 | Antagonist | 0.155 | 0.179 |
| SAF312(Libvatrep) | CC(C1=NC2=C(C(N1C3=CC=C(C#N)C=C3)=O)C=CC(O)=C2)C | Antagonist | 0.041 | -0.334 |
| Resiniferatoxin | O=C1C(C)=C[C@@]2([H])[C@@]1(O)CC(COC(CC3=CC=C(O)C(OC)=C3)=O)=C[C@@]4([H])[C@@]2(O5)[C@@H](C)C[C@]6(C(C)=C)[C@H]4OC5(CC7=CC=CC=C7)O6 | Agonist | -0.165 | -0.021 |
| PSFL2874 | O=C(N/N=C/C1=C(Cl)C(Cl)=CC=C1)C2=C(O)C=C(C=CC=C3)C3=C2 | Antagonist | 0.087 | -0.295 |

**Supplementary Table 2: Cryo-EM data collection, refinement and validation statistics**

|  | hTRPV1_Apo_  (EMDB-75616)  (PDB 11CJ) | hTRPV1_6-Iodo-CAP_  (EMDB-75617)  (PDB 11CK) | hTRPV1_Mavatrep_  (EMDB-75619)  (PDB 11CN) |
| --- | --- | --- | --- |
| **Data collection and processing** |  |  |  |
| Magnification | 165,000 | 105,000 | 105,000 |
| Voltage (kV) | 300 | 300 | 300 |
| Electron exposure (e–/Å^2^) | 50 | 50 | 50 |
| Defocus range (μm) | 1.5-2.5 | 1.5-2.5 | 0.8-2.5 |
| Pixel size (Å) | 0.73 | 0.83 | 0.83 |
| Symmetry imposed | C4 | C4 | C4 |
| Initial particle images (no.) | 1,646,090 | 1,660,569 | 2,089,583 |
| Final particle images (no.) | 153,457 | 75,798 | 198,931 |
| Map resolution (Å)  FSC threshold | 2.52  0.143 | 2.90  0.143 | 2.37  0.143 |
| Map resolution range (Å) | 2.0-2.9 | 2.2-3.2 | 2.2-3.1 |
| **Refinement** |  |  |  |
| Initial model used (PDB code) | 8GF8 | 11CN | 8GFA |
| Model resolution (Å)  FSC threshold | 2.51  0.143 | 2.90  0.143 | 2.36  0.143 |
| Map sharpening *B* factor (Å^2^) | -75.5 | -83.6 | -78.6 |
| Model composition  Non-hydrogen atoms  Protein residues  Ligands | 17,684  2,124  8 | 17,340  2,124  4 | 17,376  2,124  4 |
| *B* factors (min/max/mean Å^2^)  Protein  Ligand  Water | 1.00/120.37/35.49  12.62/108.43/47.15  N/A | 12.79/152.08/78.75  25.78/125.88/44.29  N/A | 5.39/144.74/56.58  9.41/41.73/20.08  20.7/20.7/20.7 |
| R.m.s. deviations  Bond lengths (Å)  Bond angles (°) | 0.004  0.912 | 0.004  0.904 | 0.004  0.959 |
| Validation  MolProbity score  Clashscore  Poor rotamers (%) | 1.56  5.17  0.7 | 1.47  5.50  0.54 | 1.71  9.23  0.48 |
| Ramachandran plot  Favored (%)  Allowed (%)  Disallowed (%) | 95.83  4.17  0 | 97.01  3.12  0 | 96.58  3.12  0.19 |

**Supplementary Table 2 (continued)**

|  | hTRPV1_Asivatrep_  (EMDB-75618)  (PDB 11CL) | hTRPV1_JNJ-17203212_  (EMDB-78331)  (PDB 37NU) | hTRPV1_CAP_  (EMDB-78595)  (PDB 37XE) |
| --- | --- | --- | --- |
| **Data collection and processing** |  |  |  |
| Magnification | 165,000 | 165,000 | 165,000 |
| Voltage (kV) | 300 | 300 | 300 |
| Electron exposure (e–/Å^2^) | 50 | 20 | 60 |
| Defocus range (μm) | 1.5-2.5 | 0.5-1.5 | 0.6-2.6 |
| Pixel size (Å) | 0.73 | 0.74 | 0.74 |
| Symmetry imposed | C4 | C4 | C4 |
| Initial particle images (no.) | 4,879,389 | 731,442 | 339,996 |
| Final particle images (no.) | 476,551 | 106,008 | 47,430 |
| Map resolution (Å)  FSC threshold | 2.10  0.143 | 2.34  0.143 | 3.06 |
| Map resolution range (Å) | 2.0-3.0 | 2.0-2.8 | 2.4-3.3 |
| **Refinement** |  |  |  |
| Initial model used (PDB code) | 11CN | 11CN | 11CK |
| Model resolution (Å)  FSC threshold | 2.08  0.143 | 2.34  0.143 | 3.01  0.143 |
| Map sharpening *B* factor (Å^2^) | -65.3 | -61.1 | -84.0 |
| Model composition  Non-hydrogen atoms  Protein residues  Ligands | 17,476  2,124  8 | 17,364  2,124  4 | 17,336  2,124  4 |
| *B* factors (min/max/mean Å^2^)  Protein  Ligand | 0.14/117.06/43.70  6.03/46.38/22.26 | 2.23/162.74/58.47  13.66/34.73/25.48 | 14.58/172.26/89.32  36.93/60.19/45.95 |
| Water | 17.22/29.57/21.54 | N/A | N/A |
| R.m.s. deviations  Bond lengths (Å)  Bond angles (°) | 0.004  0.866 | 0.004  0.906 | 0.005  0.984 |
| Validation  MolProbity score  Clashscore  Poor rotamers (%) | 1.22  2.86  0.21 | 1.52  6.22  1.45 | 1.56  6.14  0.59 |
| Ramachandran plot  Favored (%)  Allowed (%)  Disallowed (%) | 97.25  2.75  0 | 97.72  2.28  0 | 96.58  3.42  0 |
